# Structural basis of nucleosome remodeling by archaeal RNA polymerase during transcription elongation

**DOI:** 10.64898/2026.09.29.754952

**Authors:** Gabriele Ubartaite, Daniela Tarau, Anna Lina Bula, Winfried Hausner, Dina Grohmann, Svetlana O. Dodonova

## Abstract

Transcription occurs in the context of histone-based chromatin in eukaryotes and most archaea. The archaeal RNA polymerase and histones are ancestral to their eukaryotic counterparts, yet how RNA polymerase traverses histone-bound DNA in archaea remains poorly understood. Here, we reconstitute a nucleosome-associated transcription elongation complex (TEC) from *Pyrococcus furiosus* and capture its structure across multiple elongation states by cryo-electron microscopy (cryo-EM). High-resolution structures reveal that the RNA polymerase engages a three-dimer HPfB nucleosome positioned downstream. We identify direct physical interactions between HPfB and Rpo1N RNA polymerase subunit, establishing a defined polymerase-histone interface during transcription. Structural comparisons across defined elongation states demonstrate that RNA polymerase first translocates on DNA using the proximal histone dimer as an anchor, followed by destabilisation of the distal dimer. The DNA exiting the nucleosome is partially unwrapped and is redirected traversing the Rpo4/7 stalk. At extended transcript lengths nucleosome organisation is lost, indicating that transcription elongation ultimately disrupts histone-DNA interactions. This work provides the first structural insight into transcription through chromatin in archaea, revealing a mechanism in which the archaeal RNA polymerase actively remodels nucleosomes via DNA redirection and histone displacement, aided by direct polymerase-histone contacts.

## Introduction

In eukaryotes and most archaea, genomic DNA is packaged by histones into chromatin, the native substrate on which transcription occurs, and one that can present a major obstacle to the transcription machinery^1–4^. In eukaryotes, recent cryo-EM studies have provided detailed structural insight into how RNA polymerase II transcribes through nucleosomes and how nucleosomal DNA and histone-DNA interactions are remodelled during elongation^5–7^. These studies have established an important framework for understanding transcription through chromatin in eukaryotes. The majority of archaea also organise their genomes using histones^1,8,9^, yet how the archaeal RNA polymerase engages and traverses histone-bound DNA remains poorly understood.

Archaea are an important system for studying chromatin transcription because they share key evolutionary roots with the eukaryotic transcription machinery^10,11^ while using a distinct histone-based chromatin architecture^1,12^. Multisubunit archaeal RNA polymerases are homologous to eukaryotic RNA polymerase II, and many components of the basal archaeal transcription apparatus are ancestral to eukaryotic factors^10,11^. Archaeal histones display a conserved histone fold and are ancestral to eukaryotic histones, but typically lack the extended tails that allow regulation of eukaryotic chromatin and transcription^1,8,13,14^. Moreover, rather than forming a fixed octameric nucleosome, archaeal histones can assemble as homodimers that oligomerise along DNA through repeated histone-histone interactions, generating extended hypernucleosomes of variable length in which each histone dimer wraps approximately 30 bp of DNA^1,15^. Recent structural studies have defined the molecular basis of archaeal chromatin assembly and revealed considerable architectural diversity across archaeal histone-DNA complexes^1,16,17^. This duality of a conserved transcription machinery and divergent chromatin architecture makes archaea well-suited for identifying both universal and lineage-specific principles of transcription through chromatin.

The hyperthermophilic archaeon *Pyrococcus furiosus* (*P. furiosus*) is particularly well-suited for investigating this process. Its transcription machinery has been structurally and biochemically characterised in detail, including the transition into productive elongation and the binding of elongation factors such as Spt4/5 to the transcription elongation complex (TEC)^18–24^. Moreover, a substantial amount of genome-wide transcription data has been collected providing insights into the regulation of transcription under different environmental conditions^25,26^. *P. furiosus* encodes two canonical histone variants, HPfA and HPfB, which are highly abundant *in vivo* - a hallmark of histones in hyperthermophilic archaea^25^. In the closely related archaeon *Thermococcus kodakarensis*, histones have been shown to assemble into stable histone-DNA complexes that affect transcription elongation *in vitr*o^1,27,28^, providing a well-defined substrate for investigating how archaeal RNA polymerase navigates through histone-bound DNA.

Several lines of evidence establish that archaeal histones directly attenuate transcription^3^. Biochemical studies have shown that archaeal histones slow RNA polymerase progression and alter pausing behaviour during elongation^27–29^. At the genome-wide level, archaeal chromatin is organised around transcription start sites, with a nucleosome-free region at promoters followed by a well-positioned downstream assembly analogous to the eukaryotic +1 nucleosome^30,31^. RNA polymerase must therefore engage with and transcribe through hypernucleosomes along the gene body, raising the question of how it traverses chromatin at a molecular level - particularly given that elaborate chromatin-remodelling factors and histone chaperones that assist eukaryotic RNA polymerase II during nucleosome passage have not been identified and might be absent in archaea^3,12,32^.

Despite the overall progress in the field of archaeal transcription, the structural basis of transcription through chromatin in archaea has remained unknown. Structures are available for archaeal TECs on naked DNA^18,33^; and separately for archaeal histone-DNA assemblies^1,16,17^, but no structure has yet captured an archaeal RNA polymerase engaged with a nucleosome-like assembly. It therefore remains unclear whether direct contacts form between the polymerase and histones, how nucleosome architecture is altered during transcription, and what mechanism underlies histone displacement or retention as the polymerase advances.

Here, we reconstituted archaeal TEC-nucleosome complexes using a purified *P. furiosus* RNA polymerase, defined nucleic-acid elongation scaffolds, and histone HPfB. We initiated active transcription, allowing the RNA polymerase to encounter and transcribe beyond a positioned nucleosome. By capturing multiple transcription states using cryo-EM, we show how archaeal RNA polymerase engages a downstream HPfB nucleosome assembly, forms direct contacts with histones, translocates along the DNA while maintaining contact with histones, redirects nucleosomal DNA, and progressively destabilises histone-DNA interactions during elongation. These structures provide the first mechanistic view of an archaeal RNA polymerase transcribing through histone-bound DNA and reveal a chromatin traversal mechanism distinct from that of eukaryotic RNA polymerase II, with implications for understanding how this fundamental process evolved.

## Results

### HPfB histone forms a heterogeneous ensemble of nucleosomes and hypernucleosomes

To define the chromatin substrate used for transcription studies, we characterised the assembly of *P. furiosus* histone HPfB on DNA. Histone HPfB was chosen for further experiments due to its higher abundance in cells^25^, representing a more physiologically relevant native substrate. First, we reconstituted archaeal nucleosomal particles using HPfB (Supplementary Fig. 1A) and a 120 bp Widom601 DNA^34^. Electrophoretic mobility shift assays (EMSA) revealed a ladder-like pattern consisting of multiple discretely shifted species, consistent with binding of at least four HPfB dimers to the 120 bp DNA, with the highest-mobility-shifted species potentially corresponding to a fifth bound dimer (Fig. 1A). Next, complexes of 120 bp Widom601 DNA with HPfB were subjected to cryo-EM single-particle analysis (SPA), which revealed heterogeneous HPfB-DNA assemblies comprising both compact individual nucleosomes and larger stacked hypernucleosomes (Fig. 1B). Hereafter, we use the term nucleosome for archaeal histone-DNA complexes that wrap up to ∼120 bp of DNA, and hypernucleosome for larger assemblies formed by stacking of multiple nucleosomal units. 3D classification resolved multiple oligomeric states, including 3-dimer (hexamer), 4-dimer (octamer), and 5-dimer (decamer) assemblies (Fig. 1C), consistent with the species observed in EMSA experiments. The three structures were resolved at 3.1 Å, 2.8 Å and 3.0 Å resolutions, respectively (Table 1). In all conformations, DNA followed a left-handed superhelical trajectory around histone oligomers formed through conserved archaeal histone interfaces: stacking interface, histone dimer-dimer, and histone-DNA interfaces. Despite preservation of the canonical “closed” nucleosome architecture, histone positioning on the DNA appeared asymmetric, and portions of the histone surface were partially exposed in larger oligomers (Fig. 1C). In the 5-dimer complex, one of the peripheral histone dimers was mainly stabilized by histone-histone interactions, and only a minimal contact with the DNA (less than 10 bp engaged). This behavior also indicated that the Widom601 sequence does not position HPfB histone dimers in the same way as it does with eukaryotic histones. Overall, these data show that HPfB forms structurally heterogeneous chromatin assemblies, based on a conserved set of interfaces.

**Figure 1.**
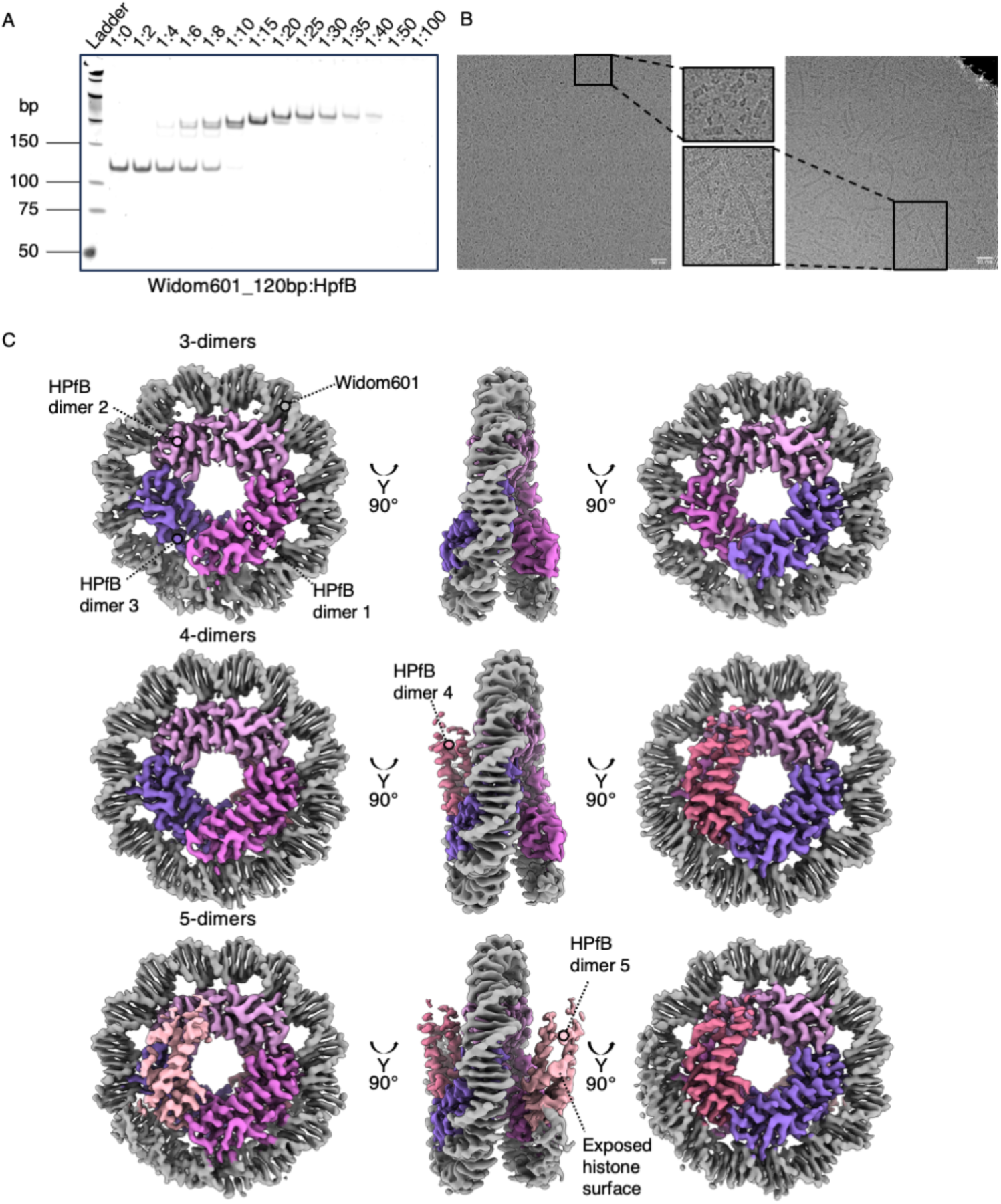
Biochemical and structural characterisation of HPfB-based nucleosome assemblies. A - EMSA showing complex formation of a 120 bp Widom601 DNA with HPfB increasing molar ratios. B - Representative cryo-EM micrographs of HPfB-Widom601_120bp complexes, with zoomed-in views highlighting minimalistic nucleosome assemblies and hypernucleosomes. Scale bars: 50 nm. C - Cryo-EM reconstructions of HPfB-Widom601_120bp assemblies containing three, four, and five HPfB dimers. Structures are shown in three orthogonal views. Individual HPfB dimers are shown in shades of pink/purple, and DNA is shown in grey.

**Table 1.**
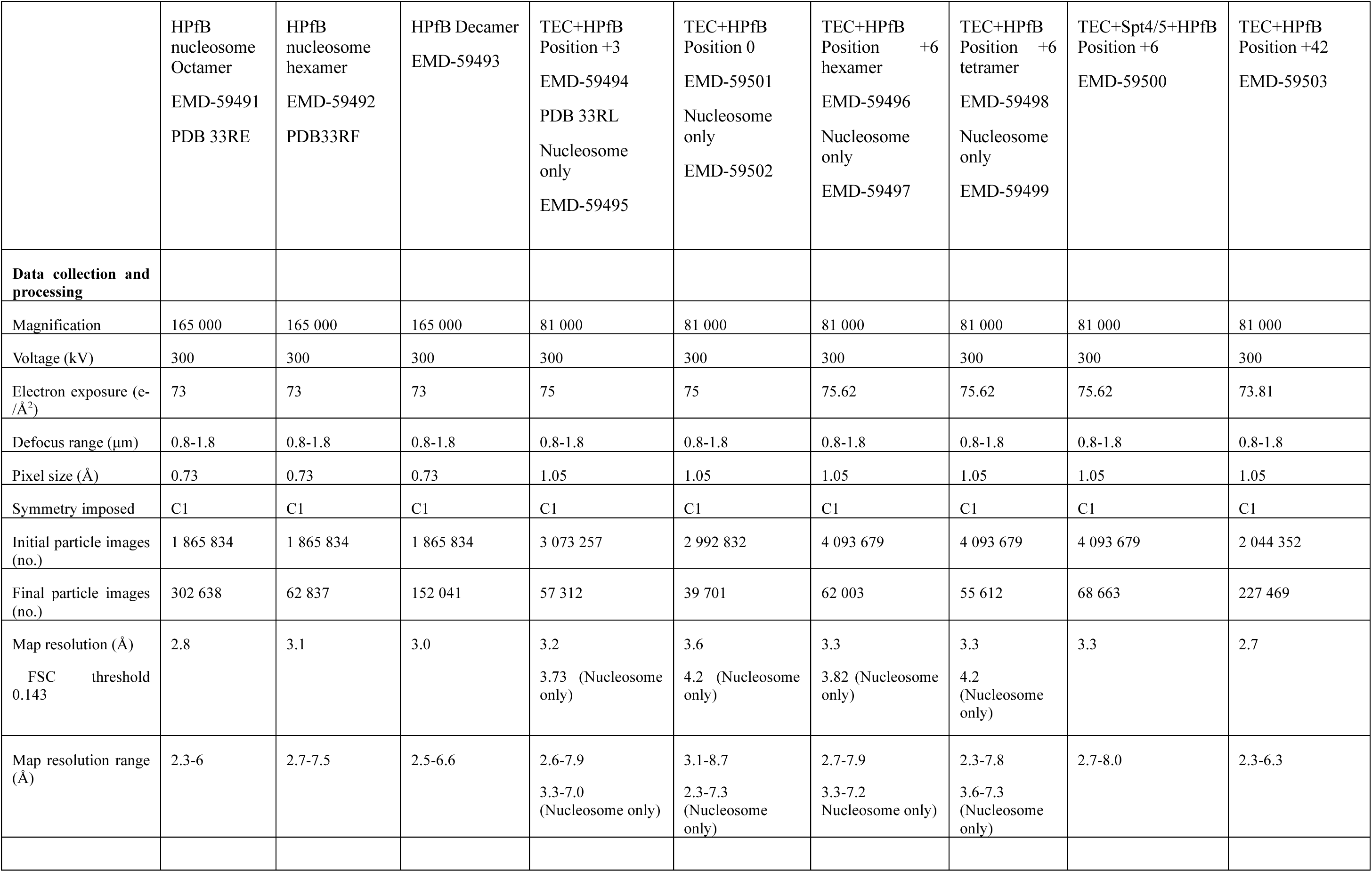
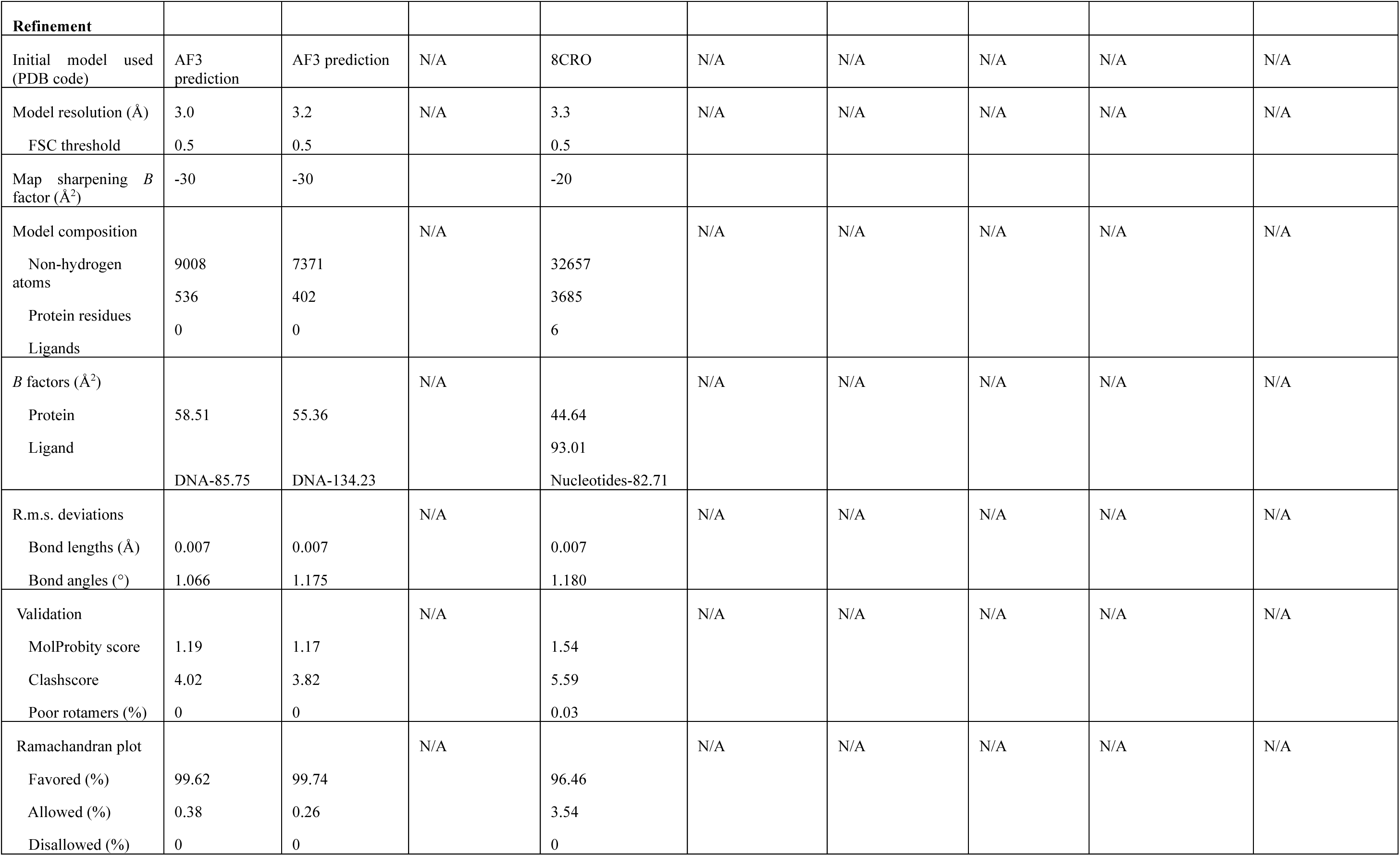

### Reconstitution and cryo-EM characterisation of a nucleosome-associated archaeal transcription elongation complex

To investigate transcription through archaeal histone-bound DNA, we reconstituted a nucleosome-associated TEC using an RNA polymerase purified from *P. furiosus* cells^18^ and recombinantly purified HPfB. The core nucleic acid elongation scaffold was based on a previously reported elongation complex^18^ and extended downstream with a Widom601^34^ DNA sequence 90bp in length to support histone assembly (Fig. 2A). The scaffold contains a transcription bubble in which a short 14 nt RNA is annealed to the template DNA strand (TS), forming a 9-bp RNA-DNA hybrid, while a stretch of the non-template DNA strand (NTS) remains single-stranded and displaced^18^. This design mimics an elongation complex approaching an archaeal nucleosome. Defined transcriptional stalling positions were introduced using CG-less and C-less cassettes. These cassettes allowed to control the position of the RNA polymerase relative to the nucleosome by varying the nucleotide set supplied during transcription, enabling preparation of homogeneous complexes for biochemical and structural analysis. Assembly of the TEC-nucleosome complex followed the workflow for TEC assembly reported previously (Supplementary Fig. 1B)^18^, followed by addition of and incubation with histones and subsequent addition of nucleotide combinations (see Methods). The assembly was confirmed biochemically: the addition of HPfB resulted in a clear mobility shift of the elongation complex in EMSA experiments (Fig. 2E; Supplementary Fig. 1C). Next, transcription assays showed that the elongation complex was fully functional and proceeded exactly until the desired stalling position at the engineered positions. Transcriptional activity in the presence of HPfB was maintained and overall comparable to the transcription pattern on free DNA (Fig. 2B-D).

**Figure 2.**
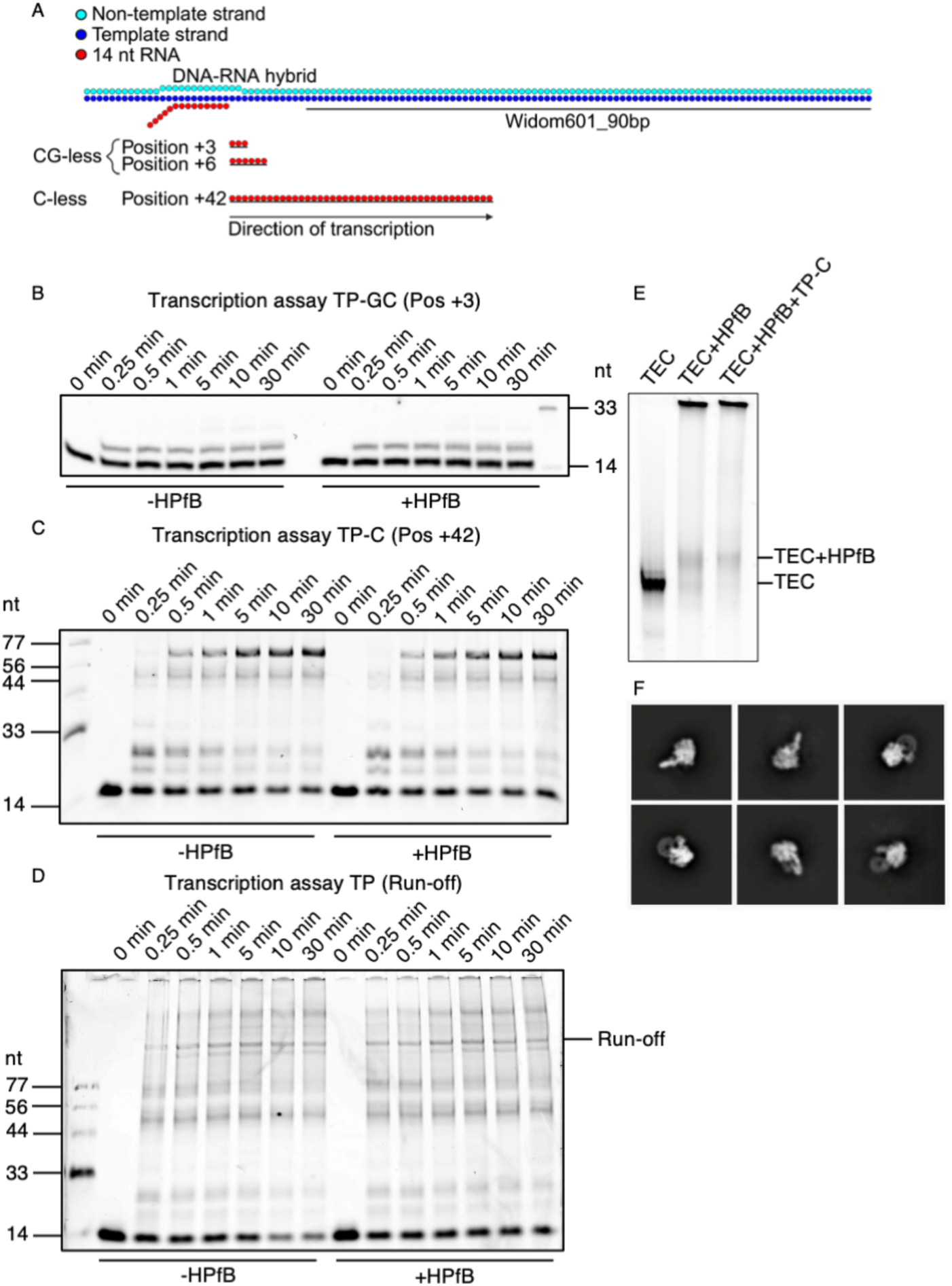
**HPfB association with the archaeal TEC does not impair transcription.**A -Schematic representation of the transcription elongation scaffold used to reconstitute the TEC-nucleosome complex. B - D Transcription elongation assays performed in absence (-HPfB) or presence (+HPfB) of histones. RNA products were analysed at the indicated time points, with increasing RNA length reflecting progressive transcription elongation. B - Transcription elongation of a TEC stalled at position +3. C - Transcription elongation from a TEC stalled at position +42. D - Run-off transcription assay. E - EMSA showing association of HPfB (pos0 and pos+42) with the TEC. F - 2D classes of TEC-nucleosome complexes.

First, complexes stalled at the end of the CG-less cassette were reconstituted. This complex is named “pos+3”, as it should elongate the RNA by three additional nucleotides compared to the initial 14 nt RNA used to assemble the TEC (“pos0”). Successful elongation of the RNA was confirmed using transcription assays (Fig. 2B, 17 nt product). The efficiency of transcription was not affected by the presence of HPfB. Assembled TEC-nucleosome pos+3 complexes were then plunge-frozen and subjected to a cryo-EM SPA workflow (see Methods). Representative 2D class averages showed well-defined RNA polymerase features, along with an additional density attributable to a nucleosome (Fig. 2F), confirming successful reconstitution of an archaeal TEC-nucleosome suitable for further structural analysis.

### Structure of the archaeal TEC-nucleosome complex reveals direct RNA polymerase-histone contacts

To define how archaeal RNA polymerase engages histone-bound DNA during transcription, we determined the structure of a TEC-nucleosome pos+3 complex by cryo-EM SPA. The reconstruction revealed a stably engaged elongation complex with a downstream nucleosome, formed by three HPfB dimers positioned directly ahead of the polymerase (Fig. 3A, Supplementary Video 1). The average resolution of the EM map was 3.2 Å, with higher local resolution within the core of polymerase reaching 2.3 Å (Supplementary Fig. 9). Although local resolution on the periphery was lower, it allowed to clearly resolve secondary structure elements, and to position the histone dimers. Additionally, a locally refined map of the nucleosome reached resolution of 3.7 Å (Supplementary Fig. 9), which allowed us to analyse the contact site between the RNA polymerase and nucleosome in detail. A combined model of the complex was built (see Methods). From the model, we conclude that the overall architecture of the RNA polymerase was consistent with previously reported archaeal elongation complexes^18^. The upstream DNA enters the central cleft of the polymerase, forming the transcription bubble, where the strands immediately separate (Supplementary Fig. 2): the NTS runs along the Rpo2 subunit, while the TS is handled by Rpo2 and Rpo1N. The TS is directed into the active site, with the arrangement of the nucleic acids consistent with a pre-translocated configuration, while the nascent RNA forms a short DNA-RNA hybrid before exiting through the RNA exit channel. Downstream of the polymerase, the DNA bends upward as it wraps around the HPfB histones.

**Figure 3.**
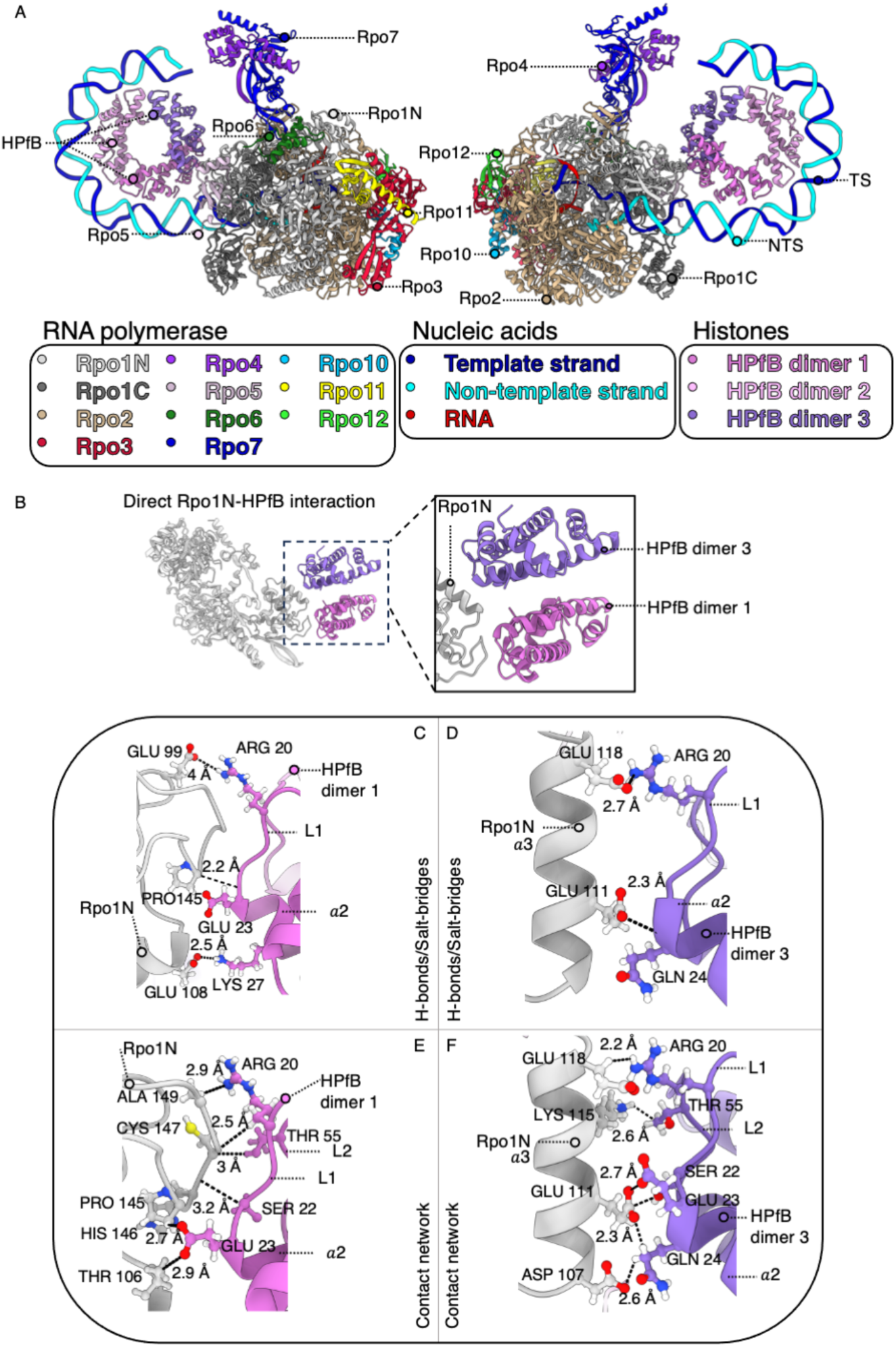
**Structure of the archaeal TEC-nucleosome complex reveals direct Rpo1N-HPfB interactions.**A - Structure of the archaeal TEC-nucleosome complex (pos+3). RNA polymerase subunits, nucleic acids, and the three HPfB dimers are coloured according to the colour scheme shown below. B - Direct interaction between Rpo1N and proximal and distal HPfB dimers. The boxed region is showing a zoom in of the interaction site, highlighting the interfaces between Rpo1N and the two HPfB dimers. C, E - Close-up view of the direct interface between Rpo1N and HPfB dimer 1, highlighting the residues involved in the interaction. D, F - Close-up view of the direct interface between Rpo1N and HPfB dimer 3. Interacting residues are shown as sticks, and distances between contacting atoms are indicated in Å. Dashed lines represent electrostatic/hydrogen bonds or contacts.

A direct interface between RNA polymerase and nucleosome was observed at the downstream DNA entry region. One interaction site was located between the Rpo1N polymerase subunit and the first (proximal) HPfB dimer, involving the histone loop L1 and part of the α2 helix (Fig. 3B, Supplementary Video 1). In this interface, HPfB residues Arg20, Glu23 and Lys27 formed electrostatic interactions or hydrogen bonds with Rpo1N residues Glu99, Pro145 (backbone) and Glu108 (Fig. 3C, 3E; Supplementary Fig. 3, Supplementary Fig. 4). A second interaction site involved the L1 and L2 loops of the third HPfB dimer and Rpo1N, centred on HPfB residues Arg20, Gln24 (backbone) and Rpo1N residues Glu118 and Glu111 (Fig. 3D, 3F; Supplementary Fig. 3, Supplementary Fig. 4). An additional contact was observed between loop L2 of the third HPfB dimer and residues 56-59 in a loop region of the Rpo5 subunit. These data reveal direct physical coupling between archaeal RNA polymerase and nucleosome during elongation and establish a defined polymerase-histone interaction interface within the TEC-nucleosome complex. Importantly, the first histone dimer is only partially engaged with the DNA, wrapping only ∼15 bp, compared to the complete wrapping of 30 bp in the isolated nucleosome complex. The rest of the histone dimer interface is positioned further away from the DNA, and instead engages with the polymerase, serving as the main contact point. Overall, ∼50 bp of DNA is directly wrapped by the complex of three histone dimers, which are further stabilized by histone-histone interactions, compared to the full 90 bp wrapped around three histone dimers in the nucleosome complex alone (Fig. 1C).

The Rpo1N residues involved in interactions with the proximal histone dimer are highly conserved within the entire order *Thermococcales*, with key residues (Supplementary Fig. 5) showing over 95% identity. Note that the Rpo1N sequence is also strongly conserved across *Thermococcales* in general, with >84% sequence identity. Histone sequences are similarly highly conserved within this group (∼90% identity). Together, these observations suggest that the polymerase-histone interaction that we report is likely relevant across the majority of archaeal species in the order *Thermococcales*. Beyond *Thermococcales*, however, the conservation of histone-interacting residues decreases.

### Structure of the archaeal TEC-nucleosome complex reveals DNA trajectory and interaction with the polymerase stalk domain

Inspection of the TEC-nucleosome complex revealed substantial distortion of the downstream nucleosomal DNA upon engagement with the RNA polymerase in comparison to a free nucleosome structure. In contrast to the continuous superhelical path observed in isolated HPfB nucleosomes, DNA within the TEC-nucleosome complex adopted a partially unwrapped configuration at the distal region of the nucleosome (Fig. 3A, Fig. 4A). The distal downstream DNA end was redirected away from the canonical nucleosomal trajectory and positioned along the RNA polymerase surface towards the Rpo4/7 stalk domain (Fig. 4A). Approximately 30 bp of distal DNA could not be reliably modelled due to reduced local resolution, indicating increased conformational flexibility within this region, but the overall DNA trajectory could be clearly recognized as it approached the polymerase stalk (Fig. 4C). Electrostatic surface analysis revealed a positively charged region directly on the stalk domain that closely matched with the observed DNA trajectory and potential DNA-stalk interaction interface (Fig. 4B). The redirected DNA remained in close proximity to this positively charged surface, consistent with electrostatic stabilisation of the altered DNA path. These observations indicate that RNA polymerase remodels archaeal nucleosome organisation during elongation by promoting partial DNA unwrapping and redirecting downstream DNA along the polymerase surface towards the stalk domain.

**Figure 4.**
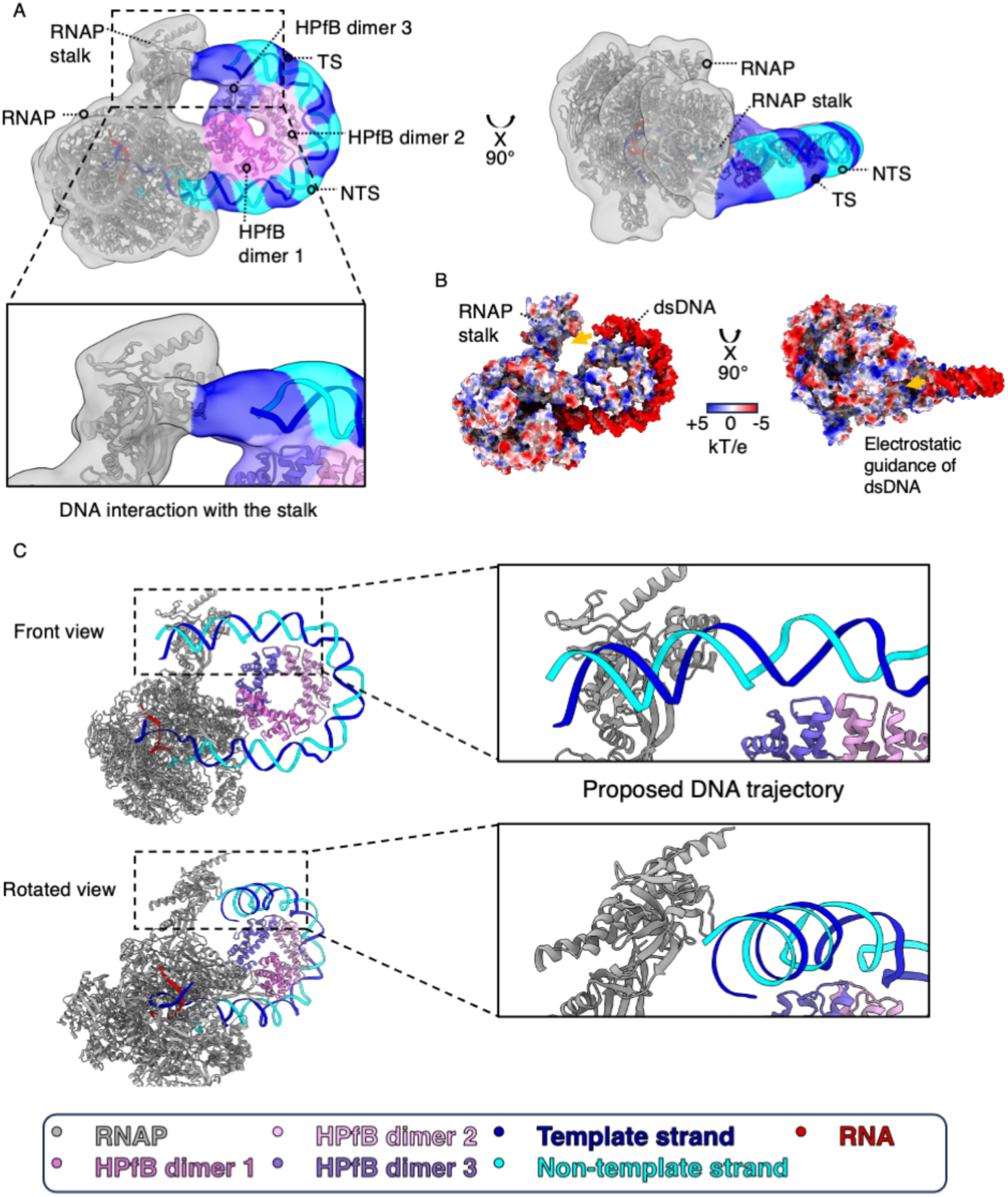
DNA redirection towards the RNA polymerase stalk. A - Archaeal TEC-nucleosome complex (pos+3) showing the interaction between nucleosomal DNA and the RNA polymerase stalk. The colour scheme for the complex is indicated below. The cryo-EM density is Gaussian-filtered and displayed at a higher contour level to highlight the DNA-stalk interaction. B - Electrostatic surface representation of the archaeal TEC-nucleosome complex highlighting the direction of nucleosomal DNA and the opposing electrostatic potentials of the DNA and RNA polymerase stalk. C - Proposed trajectory of nucleosomal DNA through the TEC-nucleosome complex, guided by the cryo-EM density, showing its redirection towards the RNA polymerase stalk.

### Cryo-EM reveals multiple intermediates of archaeal TEC transcribing through a nucleosome

To investigate how transcription progression influences nucleosome organisation, we have reconstituted several assemblies of TEC-nucleosome complexes where the RNA polymerase was located at defined positions: 0, +3, +6, and +42 (Fig. 2A) on the template DNA. After transcription, each sample was subjected to the cryo-EM SPA pipeline. The structures were resolved at overall resolutions ranging from 2.5 to 3.6 Å, with nucleosome-focused refinements ranging from 3.7 to 4.2 Å where applicable (Table 1). Across the position 0, +3, and +6 SPA datasets, the RNA polymerase remained engaged with the first histone dimer of the nucleosome, and the overall architecture of the RNA polymerase was also similar in those complexes (Fig. 5A-D, Supplementary Video 2-3). The RNA in the exit tunnel was increasingly longer (Fig. 6, 7B), consistent with our functional assays (Fig. 2 B-D). Comparison of the maps revealed lateral displacement of the downstream DNA between the pos 0, +3 and +6 states (Fig. 5F-G), in the direction orthogonal to the nucleosome plane - between the states pos 0 and +6. This shift occurred without major rearrangement of the proximal first HPfB dimers, indicating that transcription elongation into chromatinized DNA is accompanied by progressive translocation of the RNA polymerase along DNA relative to the histone core, where the Rpo1N-histone contact remains stable. The +6 dataset revealed two co-existing structural conformations (Fig. 5C-E). In one state, the nucleosome retained three intact HPfB dimers (hexamer) (Fig. 5C). In the second state, the density corresponding to the distal dimers was more fragmented and weaker than that of the 3-dimer state at the same density threshold. Therefore, we refer to this state as a 2-dimer (tetramer) state, as only the central four histones are well resolved (Fig. 5D). This region of the nucleosome was substantially rearranged, accompanied by destabilisation and loss of the third histone dimer and repositioning of adjacent DNA (Fig. 5E, 5G, Supplementary Video 2). These observations indicate that continued elongation promotes structural heterogeneity within the distal nucleosome region. To assess the consequences of extended transcription, a TEC stalled at position +42 was analysed. In contrast to earlier elongation states, no stable histone or nucleosome density could be resolved in the downstream DNA region (Fig. 6A). Although downstream DNA remained visible next to the entry site of the RNAP, it no longer adopted a nucleosome-like organisation, indicating most likely a situation in which the histone dissociated as a result of complete nucleosome destabilization.

**Figure 5.**
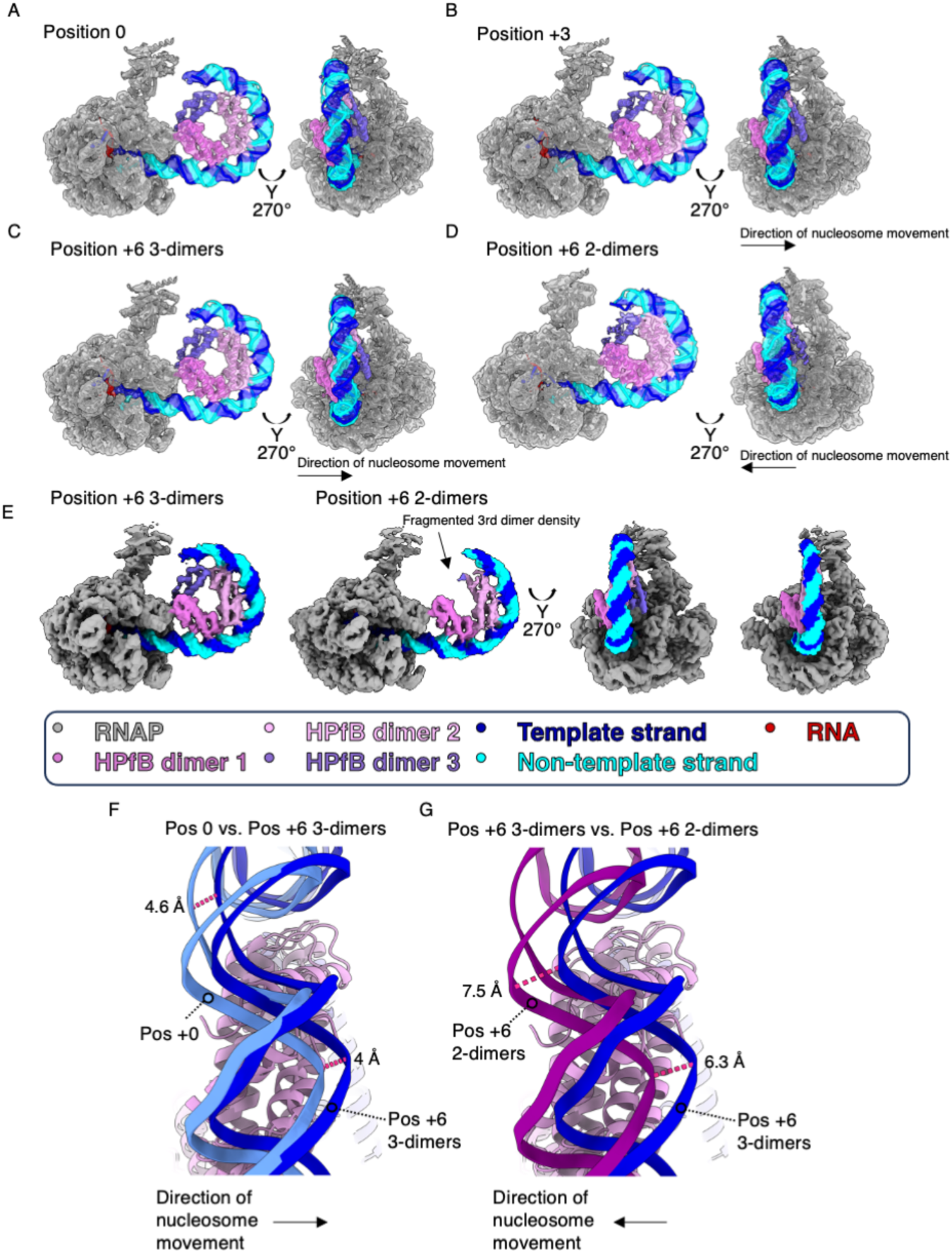
**Structural rearrangements of the TEC-nucleosome complex during transcription elongation.**A-D - Cryo-EM maps of the TEC-nucleosome complexes with fitted atomic models at positions 0, +3, +6 (3-dimer state), and +6 (2-dimer state), respectively. The direction of nucleosome movement relative to the pos0 state is indicated by arrows. E - Comparison of the pos+6 TEC-nucleosome complexes containing 3 or 2 HPfB dimers, with cryo-EM maps displayed at the same contour level. The colour scheme is indicated below. F - Comparison of nucleosomal DNA positions between the pos0 (light blue) and pos+6 (3-dimer; dark blue) states, showing DNA redirection during transcription elongation. G - Comparison of nucleosomal DNA positions between the position +6 (3-dimer; dark blue) and position +6 (2-dimer; dark purple) states, highlighting further DNA redirection associated with loss/destabilization of the third HPfB dimer. Dashed lines indicate distances between corresponding DNA positions, with distances given in Å.

**Figure 6.**
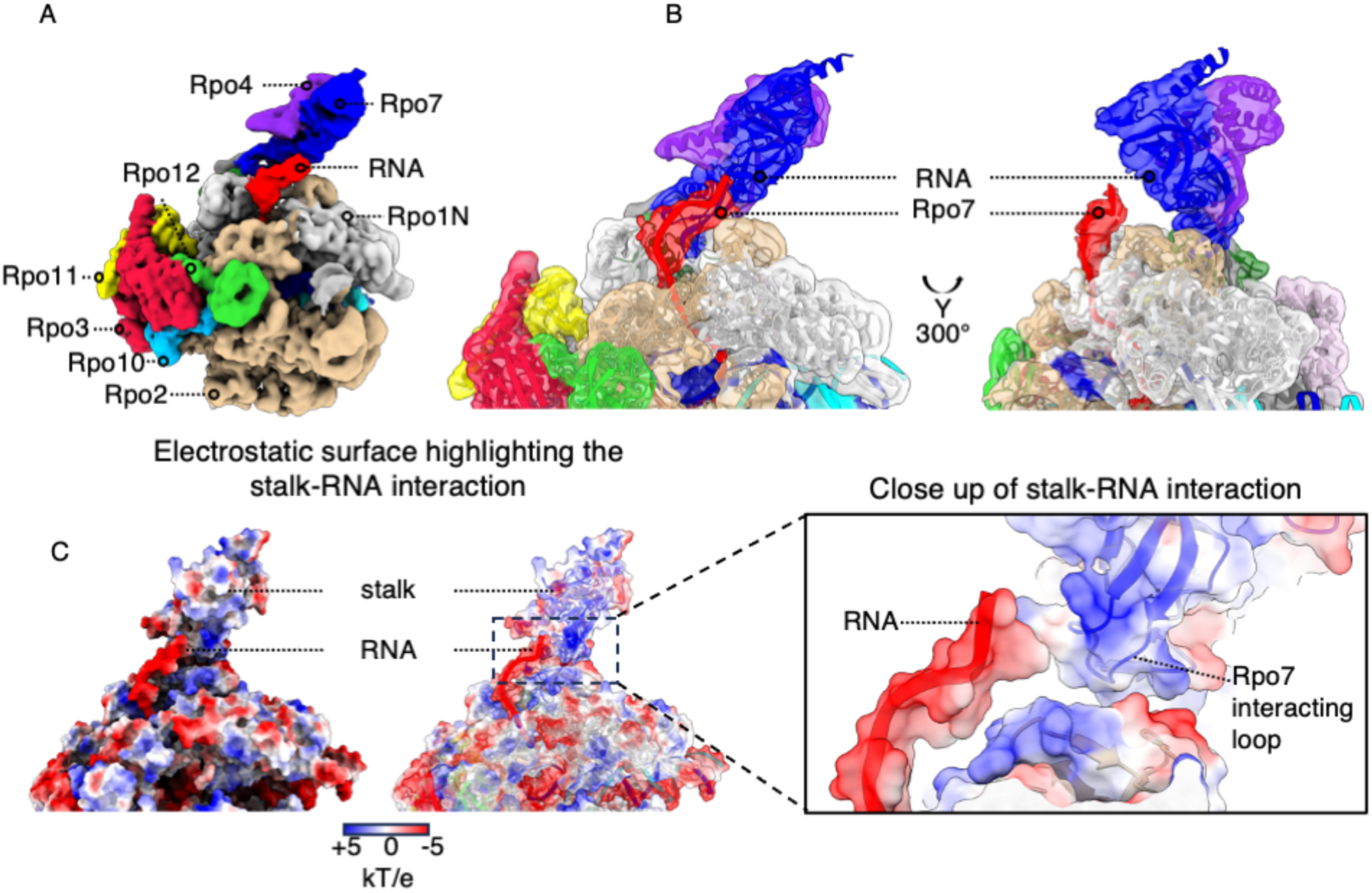
**Emerging RNA interaction with the RNA polymerase stalk.**A - Cryo-EM density map of the archaeal TEC at pos+42, with the RNA polymerase subunits and emerging RNA. No HPfB density was observed. B - Close-up views of the emerging RNA and RNA polymerase stalk, including an electrostatic surface representation of the stalk–RNA interface. The Rpo7 interacting loop is indicated. C - Electrostatic surface representation of the stalk–RNA interface showing the negatively charged emerging RNA and the positively charged surface of the Rpo7 stalk region, highlighting their complementary electrostatic interaction.

Overall, in the +3, +6 and +42 TEC-nucleosome complexes, the density corresponding to the newly synthesized RNA increased in length, confirming stepwise transcriptional progression of the complexes (Fig. 7B). These structural observations are also supported by the transcription assays (Fig. 2B-D), which showed that the main transcription products match the engineered constructs (Fig. 2). Taken together, these results indicate that histones do not fully block elongation. As expected, in the +42 complex, the RNA density was longest and extended through the RNA exit channel towards the RNA polymerase stalk (Fig. 6B). The observed RNA trajectory supports a direct interaction with a positively charged patch on the stalk^35^ (Fig. 6C). The DNA and RNA ends in the EM maps are relatively flexible, we cannot unambiguously confirm that they could bind the stalk simultaneously, although the approximate trajectory is consistent with this possibility.

**Figure 7.**
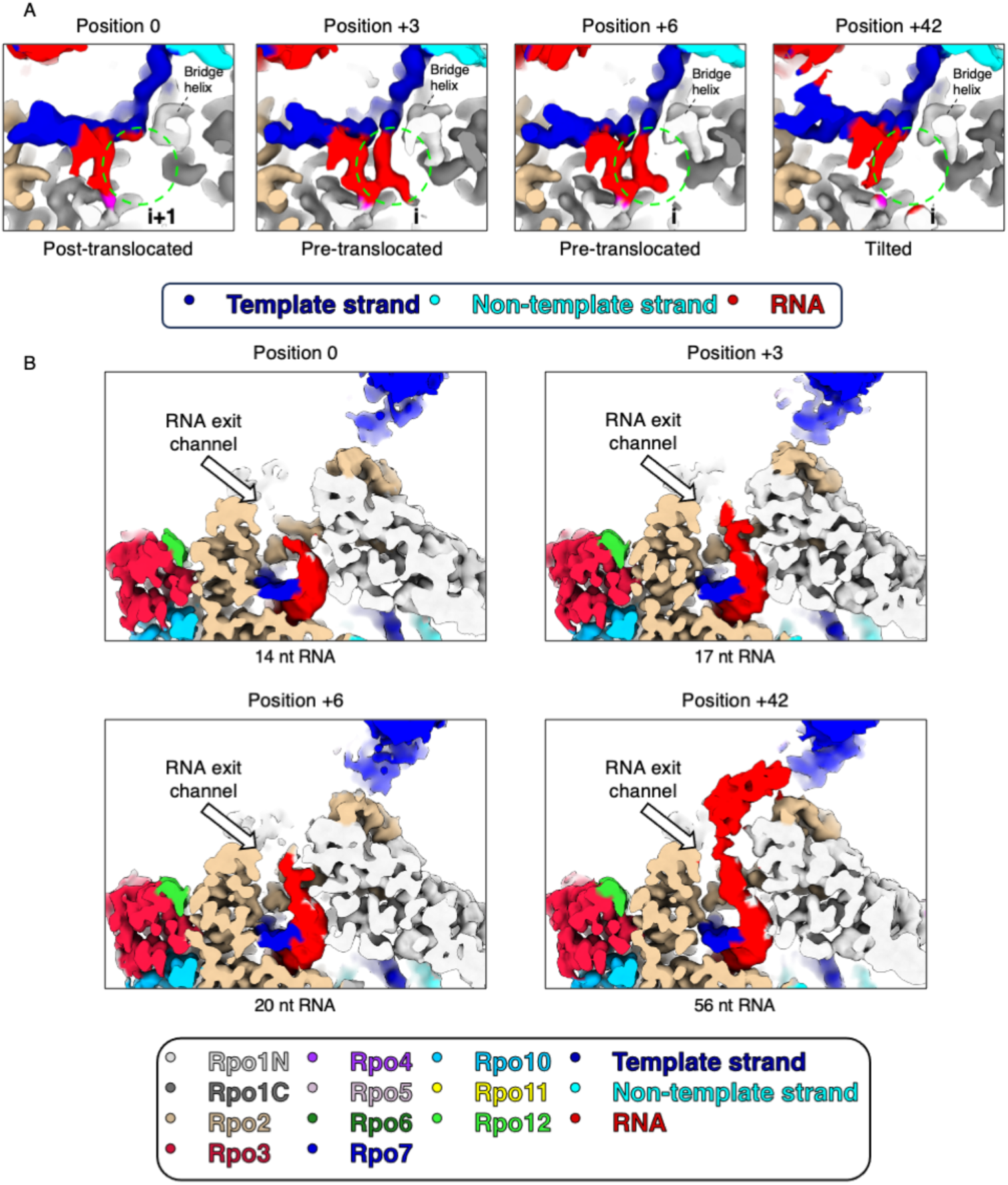
Structural changes in the active center of the RNA polymerase and RNA density in the RNA exit channel during transcription elongation. A - Cryo-EM density maps of the RNA polymerase active centre at positions 0, +3, +6 (3-dimer), and +42. The template strand, non-template strand, and RNA are coloured according to the scheme indicated below. The bridge helix and i+1 or i position are indicated, with the i+1 and i positions highlighted by a green dashed circle. Position 0 represents the post-translocated state, positions +3 and +6 represent pre-translocated states, and position +42 adopts a tilted conformation. B - Cryo-EM density maps showing the RNA exit channel at positions 0, +3, +6 (3-dimer), and +42, corresponding to RNA lengths of 14, 17, 20, and 56 nt, respectively. The emerging RNA and RNA exit channel are indicated, showing the progressive extension of RNA through the channel during transcription elongation.

**Figure 8.**
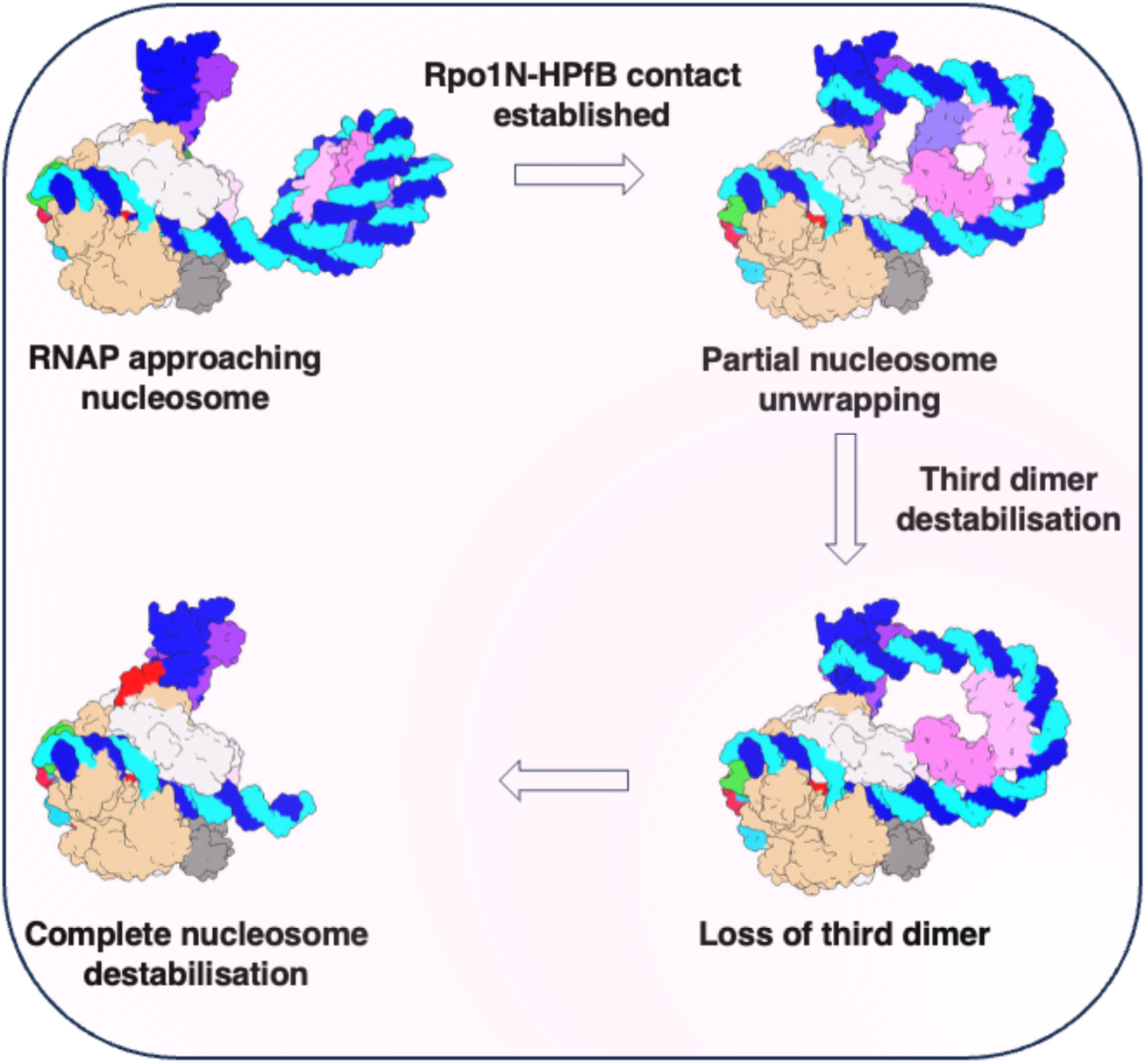
Proposed model of nucleosome remodelling by archaeal RNA polymerase during transcription elongation. The model summarizes the proposed sequence of structural rearrangements underlying nucleosome remodeling during transcription elongation. As RNA polymerase approaches the HPfB-based nucleosome, Rpo1N establishes a direct contacts with the proximal and distal HPfB dimers, accompanied by partial DNA unwrapping. Continued transcription promotes destabilisation and loss of the distal third HPfB dimer, followed by further nucleosome disruption as the polymerase progresses through the nucleosome.

Additionally, a density corresponding to the elongation factors Spt4/5 was observed in a subpopulation of particles (Supplementary Fig. 7E, Supplementary Fig. 12). Proteomic analysis of the purified RNA polymerase sample previously confirmed the presence of Spt4/5 even in the absence of nucleic acids when the RNAP was purified directly from *P. furiosus* cell mass^18^, indicating that a fraction of polymerase complexes retained this factor during sample preparation. The density occupied the canonical Spt4/5-binding site observed in previously reported archaeal elongation complexes^18^ and did not clash with the downstream nucleosome. Thus, although Spt4/5 was not a defined component of the reconstituted complex, its presence in a subset of particles indicates that it can be accommodated within the TEC-nucleosome complex.

Together, these data support a model in which transcription drives progressive nucleosome rearrangement and destabilisation during traversal of archaeal histone-bound DNA.

## Discussion

The structures presented here provide the first structural view of an archaeal RNA polymerase-nucleosome complex. Together, they reveal how the archaeal transcription machinery engages nucleosome-like assemblies and remodels them during elongation. Our data support a model in which archaeal RNA polymerase traverses chromatin through a combination of direct polymerase-histone contacts, redirection of downstream DNA, and progressive destabilisation of histone-DNA interactions.

A central finding of this study is that the archaeal RNA polymerase directly contacts the histones within the nucleosome during elongation (Fig. 3A-B). In the TEC-nucleosome structures, Rpo1N and Rpo5 interact with HPfB dimers positioned downstream of the polymerase. These contacts define interaction points between the polymerase and the histone-bound DNA substrate. In particular, the proximal HPfB dimer forms a prominent interface with Rpo1N (present in pos 0, +3, +6 complexes) and remains associated with the polymerase even when the polymerase transcribes 6 nt into the histone region (Fig. 3C, 3E, Supplementary Fig. 3). This suggests that the polymerase does not simply push against a passive histone-DNA barrier but instead engages the archaeal nucleosome through a defined protein-protein interface, and translocates along the DNA while staying engaged with the histones of the nucleosome.

This interaction provides a different perspective for previous biochemical observations which showed that archaeal histones influenced transcription *in vitro*^27–29,36^. An earlier study showed that the mutation of residue Arg20 in the histone altered transcriptional behaviour, increasing transcription rates through minimal histone-DNA complexes^27^. In our structures, Arg20 contributes not only to conserved canonical histone-DNA contacts but also to the direct interface with Rpo1N. Therefore, the effect of Arg20 mutations may reflect a combination of weakened histone-DNA binding and altered polymerase-histone engagement. However, because transcription in the presence of this stable Rpo1N-HPfB interaction was not impaired relative to free DNA in our assays, the increased transcription rates associated with Arg20 mutants are likely to arise primarily from weakened histone-DNA interactions, while the contribution from the polymerase-histone interface remains uncertain.

The direct polymerase-histone interface described here is likely to be particularly relevant within the order *Thermococcales*. The Rpo1N residues involved in HPfB binding are strongly conserved in this order, as are archaeal histones themselves (Supplementary Fig. 5). This conservation suggests that the interaction observed in our *P. furiosus* structures may represent a conserved feature of chromatin transcription within the same order. However, conservation of these residues decreases outside *Thermococcales*, indicating that the same interface may not be universally maintained across Archaea (Supplementary Fig. 5). Future comparative studies will be important to determine whether distantly related archaeal lineages use similar polymerase-histone contacts or have evolved alternative strategies for transcription through chromatin.

The mode of polymerase-nucleosome engagement observed here differs from that described for the eukaryotic RNA polymerase II (Supplementary Fig. 6B). In eukaryotic RNAPII-nucleosome complexes, the polymerase primarily interacts with nucleosomal DNA while progressing through the nucleosome^5–7^. Direct contacts between RNAPII and histones appear more limited. For example, a contact between Rpb2 and the H2A-H2B dimer has been observed when RNAPII pauses near the nucleosome Superhelical Location SHL−2^6^. More recently, a structure of RNAPII transcribing through a hexasome-octasome revealed a direct contact between the clamp of Pol II and an H3-H4 dimer^7^. The polymerase interface in that structure lies close to the archaeal Rpo1N interaction identified here, whereas the histone interface is distinct and the nucleosome adopts a markedly different orientation relative to the polymerase. Together, these observations suggest that although both archaeal and eukaryotic polymerases can establish direct contacts with histones, the molecular details of these interactions and the geometry of polymerase-nucleosome engagement differ substantially. Transcription also has distinct consequences for nucleosome architecture in archaeal and eukaryotic systems. During eukaryotic RNAPII passage, the histone octamer remains intact through coordinated DNA unwrapping and rewrapping, a process often supported by additional factors^6,37–39^. Although direct RNAPII-mediated displacement of an H2A.B-H2B dimer has also been reported^39,40^, eukaryotic nucleosome traversal generally involves a more elaborate chromatin-associated machinery^32,41^. In contrast, our structures show that archaeal RNA polymerase can engage, distort, and destabilise a histone-bound DNA substrate in the absence of additional chromatin factors. This suggests that, in archaea, chromatin remodelling during transcription can be driven primarily by the geometry and surface properties of the elongation complex itself, at least in the context of short nucleosome-like substrates.

Consistent with this idea, nucleosome engagement by the archaeal TEC is accompanied by a marked change in the trajectory of downstream DNA. In isolated HPfB-DNA complexes, DNA follows a continuous superhelical path around the histone core. In TEC-nucleosome (pos 0, +3, +6), however, downstream DNA is redirected away from the canonical nucleosomal trajectory and towards the Rpo4/7 stalk (Fig. 5A-C). Structural comparison with the isolated HPfB nucleosome indicates that maintaining the fully wrapped DNA path would be incompatible with the position of the polymerase, particularly around Rpo1N and Rpo5 (Supplementary Fig. 6A). Thus, polymerase engagement imposes structural constraints that favour DNA redirection and partial unwrapping rather than maintenance of a canonical wrapped state (Fig. 4A, 4C). This redirected downstream DNA approaches a positively charged surface on the stalk (Fig. 4B). This interaction may help stabilise the altered DNA trajectory after it leaves the histone surface. In the pos+42 complex, the nascent RNA extends towards an adjacent positively charged region of the stalk (Fig. 6A-C). This suggests an important role of the polymerase stalk in the coordination and stabilisation of the emerging RNA transcript^35^ but also nucleosomal DNA during transcription. Flexibility of the DNA and RNA ends in the EM maps make it impossible to confidently conclude whether DNA and RNA can use the stalk surface simultaneously, or if they start to compete, especially as the RNA further increases in length. This potential interplay between DNA and RNA engagement with the polymerase stalk remains to be tested in the context of longer DNA and RNA constructs.

The multiple elongation states captured in this study (Supplementary Video 3) allow us to propose a stepwise mechanism for transcription through archaeal nucleosomes. Upon initial engagement, RNA polymerase establishes contacts with HPfB dimers in the downstream nucleosome (Fig. 3A). The proximal histone dimer remains anchored to Rpo1N, while downstream DNA is redirected towards the Rpo4/7 stalk as described above (Fig. 5A-C). As elongation proceeds, the polymerase advances while the proximal polymerase-histone interface is maintained. At the same time, the distal region of the nucleosome becomes increasingly destabilised (Fig. 5E). DNA partially unwraps from the distal histone dimer, weakening histone-DNA contacts and promoting loss of this dimer. As transcription progresses, the remaining histones are displaced/evicted, as observed in the pos+42 complex (Fig. 6A), where a stable histone density is no longer resolved on the downstream DNA. This stepwise disruption suggests that archaeal nucleosome traversal proceeds through a continuum of partially remodelled states. In the first steps of elongation through nucleosomes, the nucleosomes retain a recognisable nucleosome-like architecture but already show DNA displacement and altered wrapping. Further states reveal increased heterogeneity and destabilisation of the distal histone dimer. At later stages, the ordered histone-DNA assembly is no longer maintained. These observations indicate that transcription elongation progressively weakens the histone-bound DNA substrate.

Our findings also have implications for transcription through native archaeal chromatin, which is more heterogeneous than a single defined nucleosome-like particle. Previous MNase digestion studies in hyperthermophilic euryarchaeal species have revealed broad distributions of protected DNA fragments^1,15,42^, consistent with the presence of variable hypernucleosomal assemblies. Our structures suggest how RNA polymerase can traverse short histone-bound substrates corresponding to approximately 90-120 bp of wrapped DNA, which are highly abundant in several representatives of *Thermococcales*^15,43^. In longer hypernucleosomes, however, the DNA path would be expected to create similar steric conflicts with Rpo1N and Rpo5, as discussed above (Supplementary Fig. 6A). In short hypernucleosomal substrates, these constraints are relieved by redirection of downstream DNA towards the polymerase stalk. Such redirection may be more difficult in larger hypernucleosomal arrays, where DNA is constrained by multiple additional histone-DNA contacts. We speculate that such a clash between the RNA polymerase and a modelled longer hypernucleosomal particle could be potentially resolved by a slight rotation of the hypernucleosome relative to the RNA polymerase without redirection of the DNA path. How exactly the archaeal RNA polymerase accommodates such extended hypernucleosomes therefore remains an important question for future work from both structural and functional perspectives.

Comparison with previously reported archaeal RNA polymerase complexes indicates that transcription factors from different stages of the transcription cycle are largely compatible with the TEC-nucleosome assembly. Structural superposition of the archaeal preinitiation complex containing TBP and TFB (pdb ID: 1D3U; 1AIS; matched with pdb ID:4V1N)^44–47^ reveals no steric clashes with the nucleosome-bound TEC, indicating that simultaneous occupancy is geometrically feasible (Supplementary Fig. 7C). However, because TBP and TFB function during promoter recognition and preinitiation complex assembly, they are expected to dissociate upon promoter escape before productive elongation. Whether interaction of the RNA polymerase with the first downstream nucleosome contributes to this transition remains an interesting question for future investigation (as in RNAP II^48^). Structural superposition of TFEα (pdb ID:6KF9)^49^ onto the TEC-nucleosome assembly reveals no steric clashes with the nucleosome (Supplementary Fig. 7B), indicating that its binding is geometrically compatible with the complex. However, TFEα primarily functions during transcription initiation and promoter opening and has been shown to be displaced by Spt4/5 during the transition into elongation^50,51^. Thus, while steric hindrance does not preclude TFEα binding, its presence in the nucleosome-bound elongation complex is unlikely under physiological conditions. Structural superposition of the Spt4/5-bound elongation complex^18^ (PDB 8OKI) reveals no steric clashes with the TEC-nucleosome assembly, consistent with our observation of Spt4/5 density in a subset of particles (Supplementary Fig. 7E). Similarly, the FttA (aCPSF1) pre-termination complex (PDB 9BCU)^33^ can be accommodated on the polymerase without steric interference (Supplementary Fig. 7D). However, in the FttA model, it occupies a region adjacent to the polymerase stalk, which overlaps with the nucleosomal DNA trajectory observed in our structure. Thus, although FttA itself is compatible with the TEC-nucleosome assembly, simultaneous binding of both DNA conformations would likely require an alternative trajectory for the downstream DNA.

Overall, the complexes in this study were assembled on defined DNA scaffolds and captured at engineered stalled positions. These substrates were essential for obtaining structurally homogeneous elongation intermediates. However, native archaeal chromatin contains more variable DNA sequences, different histone variants, and hypernucleosomes of diverse lengths. Future studies will therefore be needed to understand how the archaeal transcription machinery engages longer and more complex chromatin substrates.

In summary, results presented in this study provide the first structural view of archaeal RNA polymerase-nucleosome complexes, and support a model in which archaeal RNA polymerase actively remodels histone-bound DNA during transcription elongation. This process culminates in the loss of ordered nucleosome architecture as transcript length increases. These observations show how the archaeal elongation complex directly shapes chromatin organisation, revealing a mechanism of chromatin traversal distinct from that of eukaryotic RNAPII, and illustrating how transcription through chromatin has evolved.

## Acknowledgments

This work is supported by the European Union (ERC Starting Grant 3DchromArchaea, grant agreement no. 101076671, to S.O.D), and by the internal EMBL funding. D.G. gratefully acknowledges support by the Deutsche Forschungsgemeinschaft (SFB960/A7 and GR 3840/7-1) and the core funding of the University of Regensburg for this project.

We thank: Joseph Bartho and Sebastian Unger for technical assistance with microscopy; Thomas Hoffmann and EMBL IT for IT support; Quentin Durieux-Trouilleton for helpful SPA discussions; Fredrika Rajer for overall support. We thank Robert Reichelt, Thomas Hader and Simon Dechant for technical assistance to cultivate *P. furiosus* in large-scale bio-fermenters. We furthermore thank Katharina Vogl and Lena Kampf for technical assistance. We thank Andreas Schmidbauer for fruitful discussions.

Funded by the European Union. Views and opinions expressed are, however, those of the authors only and do not necessarily reflect those of the European Union or the European Research Council Executive Agency. Neither the European Union nor the granting authority can be held responsible for them.

## Author contributions

G.U. performed all biochemical preparation, reconstitution, and structural analysis presented in this study, including purification of HPfB histone, assembly and biochemical characterization of nucleosome and TEC-nucleosome complexes, cryo-EM grid preparation, and single-particle data collection and processing, under S.O.D supervision. D.T. purified the polymerase, established and shared the protocols of the elongation complex assembly and purification under the supervision of D.G.. W.H. created the *P. furiosus* encoding a tagged RNAP variant and purified archaeal RNA polymerase and trained G.U. in archaeal transcription assay methodology. A.L.B. performed additional HPfB purification. S.O.D. conceived and, together with D.G., initiated the study. S.O.D. and G.U. wrote the manuscript with input from all authors.

## Data availability

All cryo-EM maps are deposited in EMDB: EMD-59493, EMD-59495, EMD-59496, EMD-59497, EMD-59498, EMD-59499, EMD-59500, EMD-59501, EMD-59502, EMD-59503, and, with corresponding molecular models, EMD-59491 (PDB 33RE), EMD-59492 (PDB 33RF), and EMD-59494 (PDB 33RL).

## Materials and Methods

### DNA purification

DNA construct for nucleosome reconstitution was ordered from GeneArt within pMA vector. Widom601_120bp^34^ was amplified by PCR, precipitated using the standard Sodium Acetate/isopropanol procedure, and further purified using the Resource Q column (Cytiva). Another round of NaAcetate-isopropanol precipitation of the peak fractions was performed, and the DNA was dissolved in H_2_O.

Widom601_120bp sequence:

5’-GTGCCGAGGCCGCTCAATTGGTCGTAGACAGCTCTAGCACCGCTTAAACGCACGTACGCG CTGTCCCCCGCGTTTTAACCGCCAAGGGGATTACTCCCTAGTCTCCAGGCACGTGTCAGA-3’

Primers (Sigma-Aldrich): Forward 5’-GTGCCGAGGCCGCTCAATTG-3’; Reverse 5’-TCTGACACGTGCCTGGAGACT-3’

### Synthetic nucleotides

Transcription elongation complexes were reconstituted using an elongation scaffold formed with a 125 nucleotide long template (TS) and non-template (NTS) DNA strands. The NTS has a 13 nucleotide mismatch to the TS, in order to support formation of a “mismatch bubble”. A synthetic 14 nucleotide RNA oligo was complementary to the 9 nucleotides in the template DNA strand. DNA and RNA oligonucleotides were ordered from IDT.

Elongation_scaffold-Widom601_90bp:

Position 0, Position +3, Position +42 Template strand (TS) 125 nt:

5’-TCCCCTTGGCGGTTAAAACGCGGGGGACAGCGCGTACGTGCGTTTAAGCGGTGCTAGAGGTCTCTACCACCAATTCACCCCCCTCCCCACCCCAACTACTTACGCCTGGTCATTACTAGTAC TGC-3’

Mis-matched Non-template strand (mmNTS) 125 nt:

5’-GCAGTACTAGTACGAGTTGAATACGAGTAGTTGGGGTGGGGAGGGGGGTGAATTGGTGGTAGAGACCTCTAGCACCGCTTAAACGCACGTACGCGCTGTCCCCCGCGTTTTAACCGCCAA GGGGA-3’

Position +6

This TS only differs from the TS above by one nucleotide in order to make the CG-less cassette longer - 6 nucleotides instead of 3 like above. Template strand (TS) 125 nt:

5’-TCCCCTTGGCGGTTAAAACGCGGGGGACAGCGCGTACGTGCGTTTAAGCGGTGCTAGAGGTCTCTACCACCAATTCACCCCCCTCCCCACCCCAACTAATTACGCCTGGTCATTACTAGTAC TGC-3’

Mis-matched Non-template strand (mmNTS) 125 nt:

5’-GCAGTACTAGTACGAGTTGAATACGATTAGTTGGGGTGGGGAGGGGGGTGAATTGGTGGTAGAGACCTCTAGCACCGCTTAAACGCACGTACGCGCTGTCCCCCGCGTTTTAACCGCCAA GGGGA-3’

Short RNA – AUUUAGACCAGGCG

### Recombinant HPfB expression and purification

The HPfB (Uniprot ID: O59627) sequence was codon-optimised for *E. coli* and cloned into the LIC 1B plasmid from MacroLabs (pET His6 TEV LIC cloning vector 1B – Addgene #29653). Recombinant HPfB was expressed in *E. coli* BL21-CodonPlus (DE3)-RIL for 3h at 37 °C with 0.5 mM IPTG after reaching the OD600=0.6. Cells were centrifuged at 4000 xg for 25 min at 10°C and stored at -80 °C. Cells were lysed in Lysis buffer (20 mM Hepes pH 8.0, 400 mM NaCl, 5 % Glycerol, 1x PI tablet (Protease Inhibitor Cocktail, Roche), 0.1 mg/ml Lysozyme, 2.5 mM MgCl_2_, 0.01 mg/ml of DNase I) using a sonicator. Lysate was centrifuged at 27,000 x g for 20 min at 4°C. Supernatant was incubated at 75°C for 15 min in a water bath and centrifuged again at 14000 rpm for 30 min. Supernatant was incubated with 2 ml of Ni-NTA beads for 30 min at 4 °C. Beads were washed with 5 Column Volumes (CV) ml of Low salt buffer (20 mM Hepes pH 8.0, 200 mM NaCl, 5 % Glycerol), then washed with 5 CV High salt low imidazole buffer (20 mM Hepes pH 8.0, 1000 mM NaCl, 30 mM imidazole) and this was repeated for 3 times. Finally, protein was eluted with Low salt high imidazole buffer (20 mM Hepes pH 8.0, 200 mM NaCl, 500 mM imidazole) and 14 fractions (1 ml each) were collected. The protein was dialyzed in Dialysis buffer (20 mM Hepes pH 8.0, 150 mM NaCl, 1 mM DTT) and His_6_ tag cleavage with His_6_-TEV enzyme was performed overnight at room temperature (RT). The following day, the reverse His_6_-trap was done using Ni-NTA beads to remove the His_6_ tag and His_6_-TEV. The protein was loaded on a 5 ml HiTrap SP HP (Cytiva) cation exchange chromatography column using Low SP buffer (20 mM Hepes pH 8.0 150 mM NaCl) and eluted using a linear gradient over 10 CV in High SP buffer (20 mM Hepes pH 8.0, 1000 mM NaCl). Finally, protein was dialysed overnight at RT into the Final buffer (100 mM Hepes pH 8.0, 100 mM NaCl), flash frozen and kept at -80 °C.

### Purification of RNA polymerase from *Pyrococcus furiosus*

RNA polymerase was purified from the native host *Pyrococcus furiosus* and provided by Prof. Dina Grohmann. Detailed description of *Pyrococcus furiosus* growth conditions, and purification procedure as described before^18^.

### Electrophoretic mobility shift assays (EMSA)

#### Widom601_120bp with HPfB

The reactions with HPfB were set using 1.67 ng/μl of Widom601_120bp and increasing molar ratios of histones in a Reaction buffer (100 mM HEPES pH 8.0, 100 mM NaCl, 2.5 mM MgCl_2_). Samples were incubated for 3 min at 80 °C. Glycerol was added to the reactions (6% final concentration) and loaded on a 7 % EMSA gel. The gel was run in 0.5X TBE buffer at 80V for 100 min on ice. The gel was stained with SYBR gold (SYBR™ Gold, Invitrogen) and imaged with a Typhoon imager.

#### TEC with HPfB

TEC was assembled as described in the section “Elongation complex assembly for cryo-EM analysis”. After the sample was plunge frozen, 2 μl of reaction were mixed with glycerol (6% final concentration) and diluted with reaction buffer (100 mM HEPES pH 8.0, 100 mM NaCl, 2.5 mM MgCl_2_) up to 10 μl. Samples were loaded on a 5% EMSA gel and run for 1h at RT, 150V in 1x Native Laemmli buffer.

### Transcription assays

*In vitro* transcription assays were performed based on the protocol described in Tarău et al. (2024)^18^, with minor modifications. The TEC was assembled using a 5’-Cy3-labelled 14-nucleotide RNA incorporated into an elongation scaffold. To generate the TS-RNA hybrid, 2 μL of TS DNA (400 μM), 6 μL of short RNA, and 12 μL of nuclease-free water were mixed, heated to 95°C for 3 min, and then gradually cooled to room temperature using a PCR thermocycler to allow annealing. Subsequently, 12.5 μL of the TS-RNA hybrid was combined with 28 μL RNA polymerase (1.79 μg/μL) and 10 μL of 5x Cryo-EM buffer (500 mM HEPES pH 8.0, 500 mM NaCl, 12.5 mM MgCl2). The mixture was incubated at 70 °C for 10 min to promote complex formation. Thereafter, 1.5 μL of NTS (mismatched sequence) was added, followed by an additional 10-minute incubation at 70°C.

To remove excess nucleic acids, the assembled TEC was purified using a NICK size-exclusion column (Cytiva) according to the manufacturer’s instructions. DNA concentrations of collected fractions were measured using a NanoDrop spectrophotometer, and the fraction containing the highest TEC concentration was used for subsequent transcription assays.

Transcription reactions were performed as time-course experiments in the absence of histone or in the presence of HPfB at a 1:20 molar ratio. Where indicated, TEC was pre-incubated with histones for 5 min at 70°C prior to transcription initiation.

Each reaction contained 3 μL TEC, 6 μL 5× Cryo-EM buffer, and DEPC-treated water to a final volume of 26 μL. Transcription was initiated by the addition of 3 μL nucleotide mix immediately before incubation at 70°C. Nucleotide mixes used: TP-CG (0.75 mM ATP, 0.015 mM UTP); TP-C (0.75 mM GTP, 0.75 mM ATP, 0.015 mM UTP); TP (0.75 mM ATP, 0.75 mM GTP, 0.75 mM CTP, 0.75 mM UTP).

Reactions were terminated after 15 s, 30 s, 1 min, 5 min, 10 min or 30 min by addition of 1 μL 0.5 M EDTA to chelate Mg^2+^ and stop polymerase activity. Samples were denatured by adding 30 μL of formamide loading buffer (100% formamide supplemented with 0.05% bromophenol blue), followed by incubation at 95 °C for 10 min to ensure complete strand separation. The resulting elongation products were resolved on a denaturing polyacrylamide gel containing 8 M urea and 20% acrylamide. Electrophoresis was performed at a fixed power of 20 W. A volume of 15 μL of each denatured sample was loaded per well. Fluorescent detection of Cy3-labeled products was carried out using a Typhoon Imager.

### Elongation complex assembly for cryo-EM analysis

#### Widom601_120bp with HPfB

100 ng/μl of Widom601_120bp was incubated with 20x molar excess of HPfB for 3 min at 80°C in 100 mM HEPES pH 8.0, 100 mM NaCl, 2.5 mM MgCl_2_. UltrAufoil R2/2 Au 200 grids (Quantifoil) were glow-discharged using PELCO easiGlow (Ted Pella) for 70s (0.26mB, 25mA). 3 μl of sample were applied to the grid, followed by blotting, and plunge-freezing into liquid ethane (Vitrobot Mark IV; Thermo Fisher Scientific; 100% humidity; 20 °C; 3 s blotting time, 0 blotting force).

#### TEC+HPfB

TECs were generated by initially annealing 10 μl short RNA (400 μM) with 2 μl of TS DNA (400 μM) at a 5:1 molar ratio in a total reaction volume of 20 μl. The mixture was heated to 95 °C for 3 min and then gradually cooled to allow hybrid formation. Subsequently, 20 μl of 5X CryoEM buffer, 56 μl of Polymerase (1.79 mg/ml) were added and incubated for 10 min at 70 °C in PCR cycler. 2.5 μl of NTS were added and incubated for 10 min at 70 °C. Excess of nucleotides was removed by gel filtration chromatography using the Superose 6 Increase 3.2/300 column (Cytiva) in CryoEM buffer (100 mM HEPES/KOH pH 8.0, 100 mM NaCl, 2.5 mM MgCl_2_). The concentration of DNA was estimated from peak fractions, and 20x molar excess of HPfB was added. Measured DNA concentration varied at different preparations and was ∼100-130 ng/μl. The complex was incubated for 5 min at 70 °C. For pos +3 and pos+6 sample TP-CG and pos+42 for TP-C were added accordingly and incubated for 10min at 70 °C (stock and final concentrations of NTPs were as follows: TP-CG mix (25 mM ATP, 0.2 mM UTP stock; final concentration in the reaction 1 mM ATP, 8 μM UTP); TP-C mix (25 mM ATP, 25 mM GTP, 0.5 mM UTP stock; final concentration 1 mM ATP, 1 mM GTP, 20 μM UTP). Samples were immediately applied to glow-discharged grids, blotted, and plunge-frozen into liquid ethane (Vitrobot Mark IV; Thermo Fisher Scientific; 100% humidity; 20 °C; 3 s blotting time, 0 blotting force). The remaining samples were checked on EMSA gels.

### Data acquisition and processing

#### *HPfB+Widom601_120bp* (Supplementary Fig. 14)

Data were collected on a Titan Krios G4 microscope at 300 keV equipped with a Falcon4i direct electron detector (Thermo Fisher Scientific) and a Selectris X Energy filter. Serial EM^52^ software was used for acquisition. Defocus values ranged from -0.8 to -1.8 μm at a nominal magnification of 165 000x and a pixel size of 0.73 Å. The energy filter slit width was set to 10 eV. The total electron dose of 73 e/A2 was applied, and data were collected in electron counting mode. In total, 12 216 movies were collected. Data were processed using CryoSPARC 4.1.1^53^. The movie frames were aligned and dose-weighted using Patch Motion Correction^54^ and Patch CTF estimation. 11 661 micrographs were used for further processing. Particles were first picked using Blob picker, and an initial 2D classification was run. The best classes were used for the 2D Template picker. The best classes after the 2nd classification were used for Topaz^55^ training (3 times) and picking, resulting in 1 035 k selected particles. Next, we performed ab initio reconstruction into five classes, yielding one high-quality class (772 k particles) and four classes containing low-quality or contaminating particles. The best class was then used for heterogeneous refinement. After a few rounds of 3D classifications and heterogeneous refinements, we could find hexameric (62.8 k particles, 3.1 Å), octameric (302.6 k particles, 2.8 Å), and decameric assemblies (152 k particles, 3.0 Å). The best models were refined using NU-refinement, reference motion corrected, globally CTF refined and locally refined. The final resolutions were estimated using the 3D validation procedure in CryoSPARC^53^.

#### TEC+HPfB data collections

Cryo-EM data collection parameters are summarized in Table 1. Data were processed using CryoSPARC^53^. The movie frames were aligned and dose-weighted using Patch Motion Correction^54^ and Patch CTF estimation. Particles were first picked using the Blob picker. The best classes were used for ab initio generation and used for 3D classification. Once the class with nucleosome density was detected, a focused nucleosome mask was generated and all particles were 3D classified with the focused mask. An additional round of 3D classification was performed to get rid of bad particles or investigate different conformations. The best maps were then NU-refined, reference-motion corrected, global CTF was refined, followed by local refinement and map sharpening.

#### *TEC+HPfB Position 0* (Supplementary Fig. 8)

Particle picking was performed using the blob picker algorithm with following parameters: 150-250 A; 1 separation distance, yielding a total of 2 992 832 particles. Particles were extracted with a box size of 430 pixels. These particles were subjected to two consecutive rounds of 2D classification to remove non-particle images and poorly aligned classes. Following this, 941 602 particles were maintained for further analysis. An initial 3D model was generated using ab initio reconstruction, followed by NU-refinement to improve map quality. All particles were subsequently re-centered by applying new coordinates (215, 215, 165), and focused 3D classification was performed using a mask encompassing the region corresponding to the extra density. One class containing 248 983 particles exhibited a well-defined histone dimer and nucleosome-like density. These particles were subjected to NU refinement and an additional round of focused 3D classification. The best-resolved class from this step contained 72 251 particles. A further round of NU refinement and focused 3D classification was conducted, resulting in a final high-quality class containing 39 701 particles. The final reconstructed map reached an overall resolution of 3.6 Å.

To assess the nucleosome density, a particle subtraction procedure was performed using the final map. Following density reconstruction, local refinement was carried out to further improve map features. The final nucleosome-only reconstructed map reached an overall resolution of 4.2 Å. The final resolution was estimated using the 3D validation procedure in CryoSPARC^53^.

#### *TEC+HPfB Position +3* (Supplementary Fig. 9)

Particle picking was performed using the blob picker algorithm with the following parameters: 150-250 A; 1 separation distance, resulting in 3 873 513 particles. Particles were extracted with a box size of 460 pixels and Fourier-cropped to 230 pixels, yielding 3 073 257 particles for further processing. These particles were subjected to two rounds of 2D classification. Following 2D classification, 1 680 151 particles were retained for subsequent analysis. Three ab initio models were generated, and heterogeneous refinement was carried out to separate polymerase, polymerase dimers, nucleosome-only particles, and poorly resolved classes. The polymerase class contained 1 070 280 particles, which were further refined using NU refinement. A first round of focused 3D classification targeting additional density was performed into two classes (additional density was first detected by 3D classification without a focus mask). The best-resolved class, containing 472 223 particles, was selected for a second round of nucleosome-focused 3D classification into three classes. From these, 124 148 particles were selected, re-extracted with a 460 pixel box size, and subjected to NU refinement. Subsequently, these particles underwent an additional round of nucleosome-focused 3D classification to two classes, resulting in a final high-quality class containing 57 312 particles. The final reconstructed map achieved an overall resolution of 3.2 Å.

To assess the nucleosome density, the same particle subtraction procedure was done as for Pos 0 dataset. The final nucleosome-only reconstructed map reached an overall resolution of 3.7 Å.

#### *TEC+HPfB Position +6* (Supplementary Fig. 10, Supplementary Fig. 11, Supplementary Fig. 12)

Particle picking was performed using the blob picker algorithm with following parameters: 150-250 A; 0.9 separation distance, yielding a total of 4 093 679 extracted particles with a box size of 460 pixels, binned to 230 pixels. These particles were subjected to two consecutive rounds of 2D classification to remove non-particle images and poorly aligned classes. Following this, 982 045 particles were maintained for further analysis. An initial 3D model was generated using ab initio reconstruction, followed by NU-refinement. All particles were subsequently re-centered by applying new coordinates (215, 215, 165), and focused 3D classification into 5 classes was performed using a mask encompassing the region corresponding to the extra density (additional density was first detected by 3D classification without a focus mask). One class containing 118 434 particles exhibited a well-defined nucleosome-like density and particles were unbinned. These particles (118 106 left after re-extraction) were subjected to NU-refinement and an additional round of focused 3D classification. This resulted into two best-resolved classes containing 62 209 and 55 897 particles. Both final reconstructed maps reached an overall resolution of 3.3 Å.

To assess the nucleosome density, the same particle subtraction procedure was done as for Pos 0 dataset. The final nucleosome-only reconstructed maps reached an overall resolution of 3.8 for hexamer and 4.2 Å for tetramer.

In addition, 3D classification of 982 045 particles was conducted into 3 classes. One class revealed extra density on polymerase, which was used to generate a focused mask around that region. Focused 3D classification was performed into 5 classes. 214 735 particles from this class were NU-refined and density was identified as Spt4/5. Particles were then subjected to 3D focus classification around the nucleosomal density into 3 classes, which resulted in one class containing 68 663 particles with two histone dimers bound. Final reconstructed map reached an overall resolution of 3.3 Å.

#### *TEC+HPfB Position +42* (Supplementary Fig. 13)

Particle picking was performed using the blob picker algorithm with the following parameters: 150-250 A; 1 separation distance, yielding a total of 2,044,352 extracted particles with a box size of 460 pixels. These particles were subjected to two consecutive rounds of 2D classification to remove non-particle images and poorly aligned classes. Following this, 849 609 particles were maintained for further analysis. An initial 3D model was generated using ab initio reconstruction, followed by NU-refinement to improve map quality. All particles were reference motion corrected, NU-refined, Global CTF refined; NU-refined and Local refined leading to a map of 2.5 A. These particles were 3D classified into 10 classes. 5 classes containing good densities were NU-refined. No extra nucleosome density was observed.

RNA exit channel: A spherical mask was first generated and used for a focused 3D classification around RNA exit channel. After extra density was observed, a more focused mask was generated, and 3D classification was performed again. This resulted in a good class containing 227 469 particles. The density was NU-refined and locally refined. The final reconstructed map achieved an overall resolution of 2.7 Å.

The data was processed using EMBL Heidelberg HPC Cluster^56^.

### Structural model building and refinement

#### HPfB-Widom601_120bp

The AlphaFold 3^57^ prediction models were used for rigid-body fitting into the density using UCSF ChimeraX v1.10.1^58,59^. The model was fit into the density by ISOLDE relaxation^60^. Real-space refinement was performed in Phenix 2.0^61^. Coot v0.9.8.93^62^ was used to build in the RNA sequence and fix rotamer outliers and side chain conformations.

#### TEC+HPfB Position +3

The 8CRO atomic model for RNA polymerase^18^ and AlphaFold2^63^-predicted HPfB dimers were used for rigid-body fitting. An AlphaFold3^57^ model including the DNA sequence was generated, and the DNA segment with curvature matching the density was selected for rigid-body fitting using UCSF ChimeraX v1.10.1^59^. The model was further adjusted into the density using ISOLDE^60^. Real-space refinement was performed in Phenix 2.0^61^. Coot v0.9.8.93^62^ was used to build the RNA and correct rotamer outliers and optimize side-chain conformations. Model validation was performed using MolProbity^64,65^. Other models (pos 0, pos+6 and pos+42 were prepared by rigid body fitting all chains separately from the pos+3 model).

### MSA analysis

Sequences of archaeal Rpo1N RNA polymerase subunits annotated in UniProt^66^ were pooled and subjected to multiple sequence alignment using Clustal Omega^67^. For the overall Rpo1N MSA, 605 complete, non-fragmented, annotated sequences were used. For the *Thermococcales*-specific Rpo1N analysis, 44 sequences were used. Jalview^68^ was used for MSA visualization and analysis.

