## Supplementary Figures for "Structural basis of nucleosome remodeling by archaeal RNA polymerase during transcription elongation"

This PDF file includes:
- Supplementary Figures 1 to 14

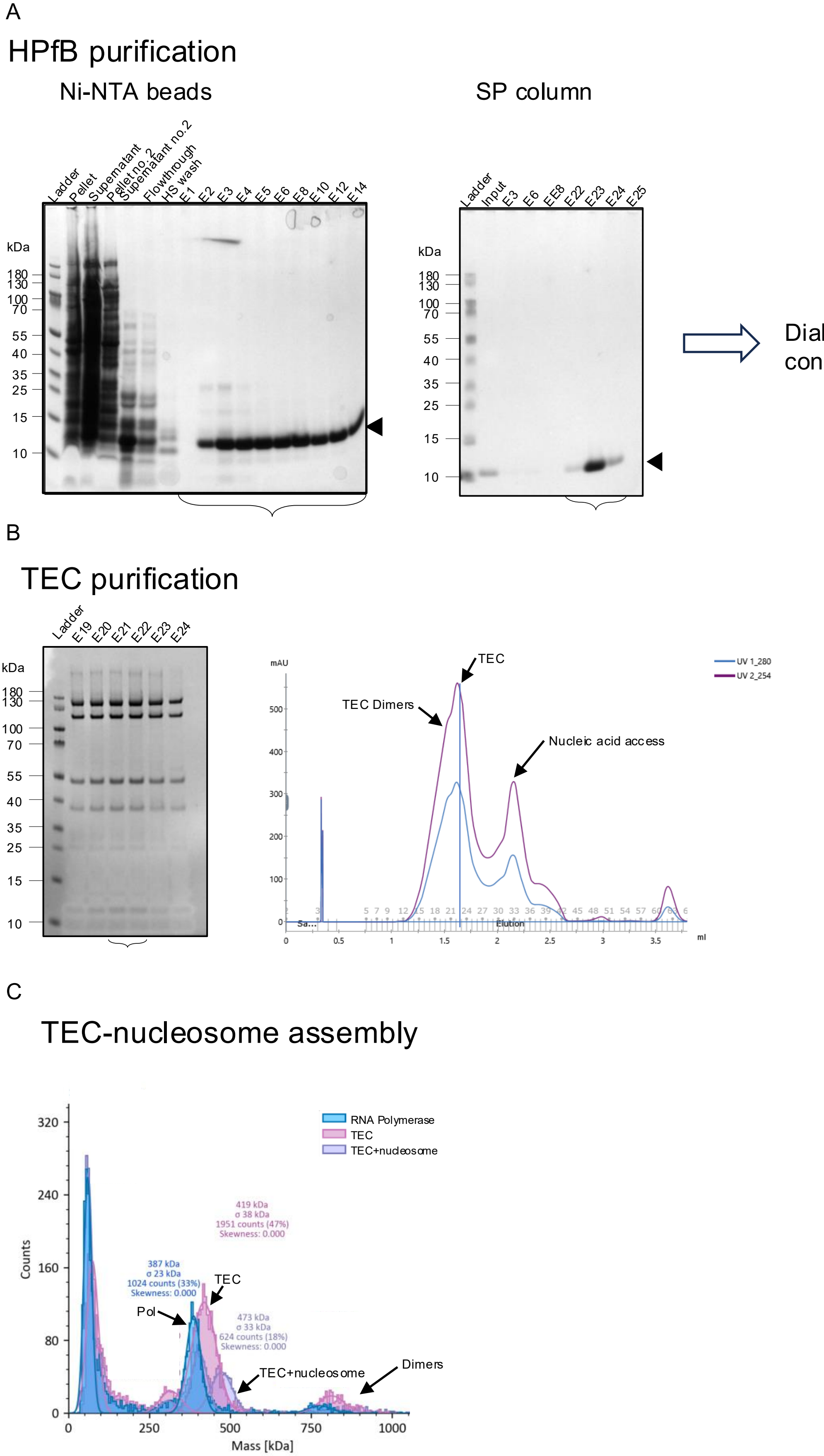


**Supplementary Figure 1. Purification and assembly of the TEC-nucleosome complex.** A - Purification of HPfB by Ni-NTA affinity chromatography followed by SP ion-exchange chromatography. SDS-PAGE analysis of the indicated fractions is shown. Fractions containing HPfB are indicated, followed by dialysis and concentration. B - Purification of the TEC. SDS-PAGE analysis of the indicated fractions and size-exclusion chromatogram is shown. The major peak corresponding to the TEC is indicated, together with the TEC dimer and nucleic acid-containing fractions. C - Assembly of the TEC-nucleosome complex analysed by mass photometry. Mass distributions corresponding to RNA polymerase, TEC, TEC-nucleosome, and dimers are indicated.

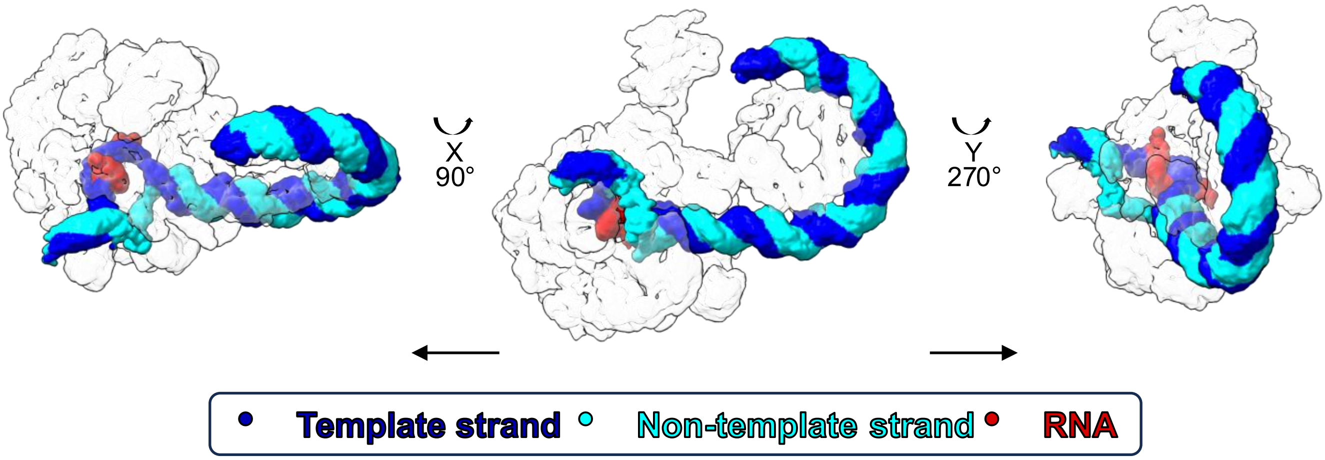


**Supplementary Figure 2. DNA-RNA pathway in the TEC-nucleosome complex at pos +3.** Three views of the TEC-nucleosome complex at position +3 showing the DNA-RNA pathway through the complex. The displayed density is a segmentation map. The template strand, non-template strand, and RNA are coloured according to the scheme indicated below. The central view is rotated by 90° and 270° around the indicated axes.

| HPfB chain | HPfB residue | Rpo1N residue | Arpeggio contact type(s) | Min. distance (Å) |
|---|---|---|---|---|
| **G** | Thr55 | His146 | proximal | 4.22 |
| **G** | Thr55 | Cys147 | **polar**, proximal | 3.15 |
| **G** | Thr55 | Gly148 | proximal | 4.81 |
| **G** | Lys57 | His146 | proximal | 4.76 |
| **I** | Gln19 | Gly148 | proximal | 4.54 |
| **I** | Arg20 | Glu99 | **ionic**, proximal | 3.95 |
| **I** | Arg20 | Lys115 | proximal | 4.51 |
| **I** | Arg20 | Cys147 | weak_polar, proximal | 3.40 |
| **I** | Arg20 | Gly148 | proximal | 3.65 |
| **I** | Arg20 | Ala149 | proximal | 3.77 |
| **I** | Arg20 | Pro150 | proximal | 4.39 |
| **I** | Val21 | Cys144 | proximal | 4.60 |
| **I** | Val21 | Pro145 | proximal | 4.90 |
| **I** | Val21 | Cys147 | proximal | 4.50 |
| **I** | Val21 | Gly148 | proximal | 3.67 |
| **I** | Ser22 | Pro145 | proximal | 3.64 |
| **I** | Ser22 | His146 | **vdw, weak_polar**, proximal | 3.23 |
| **I** | Ser22 | Cys147 | proximal | 4.89 |
| **I** | Glu23 | Thr106 | weak_polar, vdw_clash, proximal | 2.88 |
| **I** | Glu23 | Arg141 | proximal | 4.31 |
| **I** | Glu23 | Val143 | proximal | 4.47 |
| **I** | Glu23 | Pro145 | **hydrophobic, polar**, vdw_clash, proximal | 3.02 |
| **I** | Gln24 | Asp107 | proximal | 4.36 |
| **I** | Gln24 | Pro145 | proximal | 4.37 |
| **I** | Lys27 | Thr106 | proximal | 4.88 |
| **I** | Lys27 | Glu108 | **hbond, ionic, polar**, vdw_clash, proximal | 2.52 |
| **O** | Arg20 | Glu118 | **ionic, polar**, vdw_clash, proximal | 2.72 |
| **O** | Arg20 | Leu119 | proximal | 3.77 |
| O | Arg20 | Lys115 | proximal | 4.51 |
| **O** | Ser22 | Glu111 | **polar, weak_polar, vdw**, vdw_clash, proximal | 2.60 |
| **O** | Glu23 | Glu111 | vdw_clash, proximal | 2.72 |
| **O** | Glu23 | Met114 | proximal | 4.30 |
| **O** | Gln24 | Asp107 | **polar, weak_polar, vdw**, proximal | 3.25 |
| **O** | Gln24 | Glu111 | **hydrophobic, polar, weak_polar, vdw**, vdw_clash, proximal | 3.08 |
| **Q** | Thr55 | Lys115 | **hydrophobic**, vdw_clash, proximal | 3.23 |
| **Q** | Lys57 | Glu108 | proximal | 4.69 |
| **Q** | Lys57 | Glu112 | proximal | 4.93 |

**Supplementary Figure 3. Arpeggio analysis of HPfB-Rpo1N interactions.** Arpeggio analysis of residues at the HPfB-Rpo1N interface. The HPfB chain and residue, corresponding Rpo1N residue, contact types, and minimum distance between residues are indicated. Bold text highlights specific interaction types identified at the interface, including hydrogen bonds, ionic, polar, weak-polar, hydrophobic, van der Waals (vdw), and van der Waals clash (vdw_clash) interactions. Proximal contacts are also reported but are not highlighted. For each residue pair, only the closest interatomic contact is reported, together with the corresponding Arpeggio contact types. Minimum distances are given in Å.

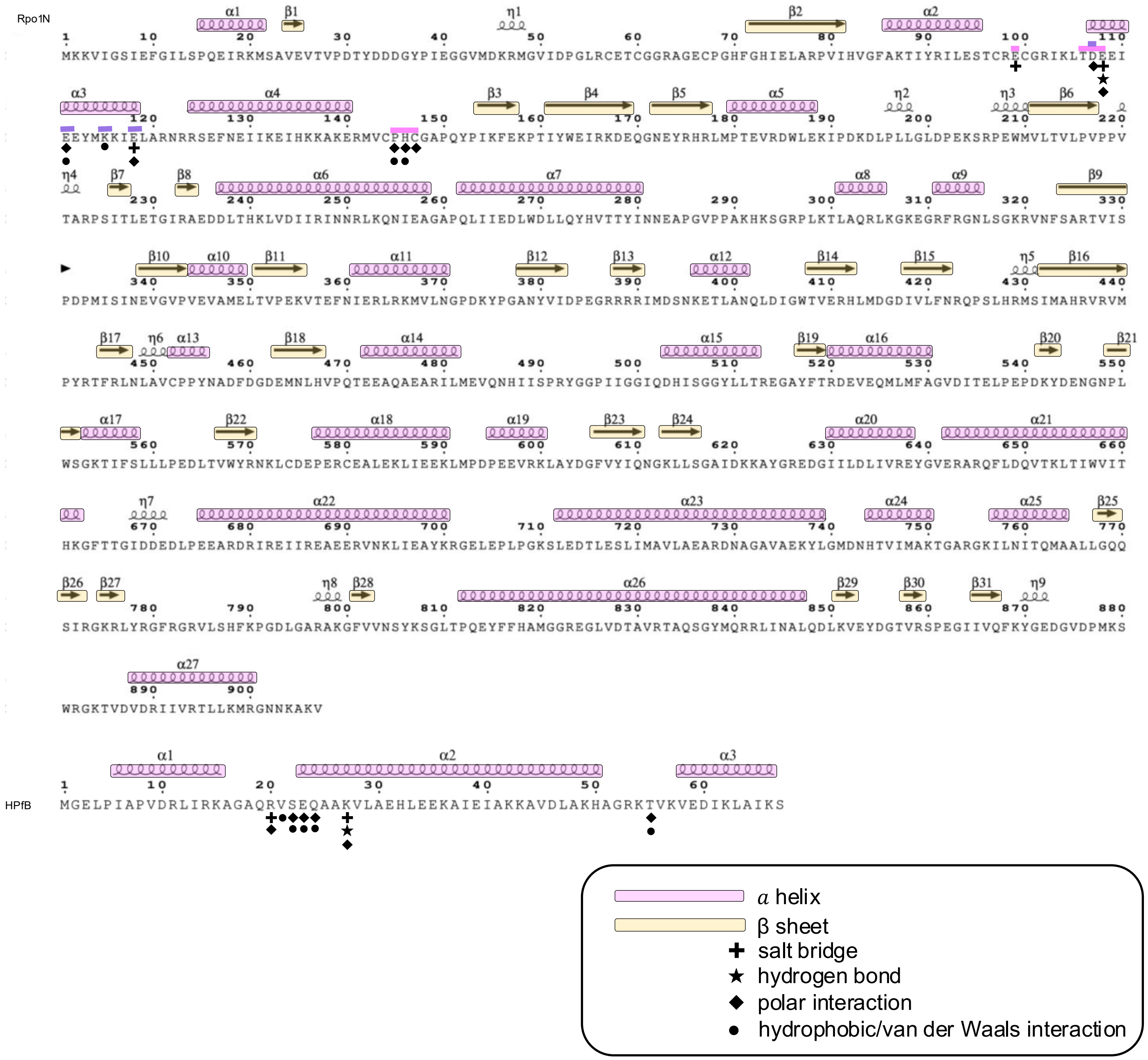


**Supplementary Figure 4. Secondary structure and Rpo1N-HPfB interaction sites.** Amino acid sequences of Rpo1N and HPfB with secondary-structure elements indicated above the sequences. Residues contacting the proximal HPfB dimer are highlighted in orchid, whereas residues contacting the distal HPfB dimer are highlighted in purple. Interaction types are labelled as follows: salt bridges +, hydrogen bonds ★, polar interactions ◆, and hydrophobic/van der Waals interactions ●. The contacting residues are not labelled. The colour scheme for secondary-structure elements is indicated.

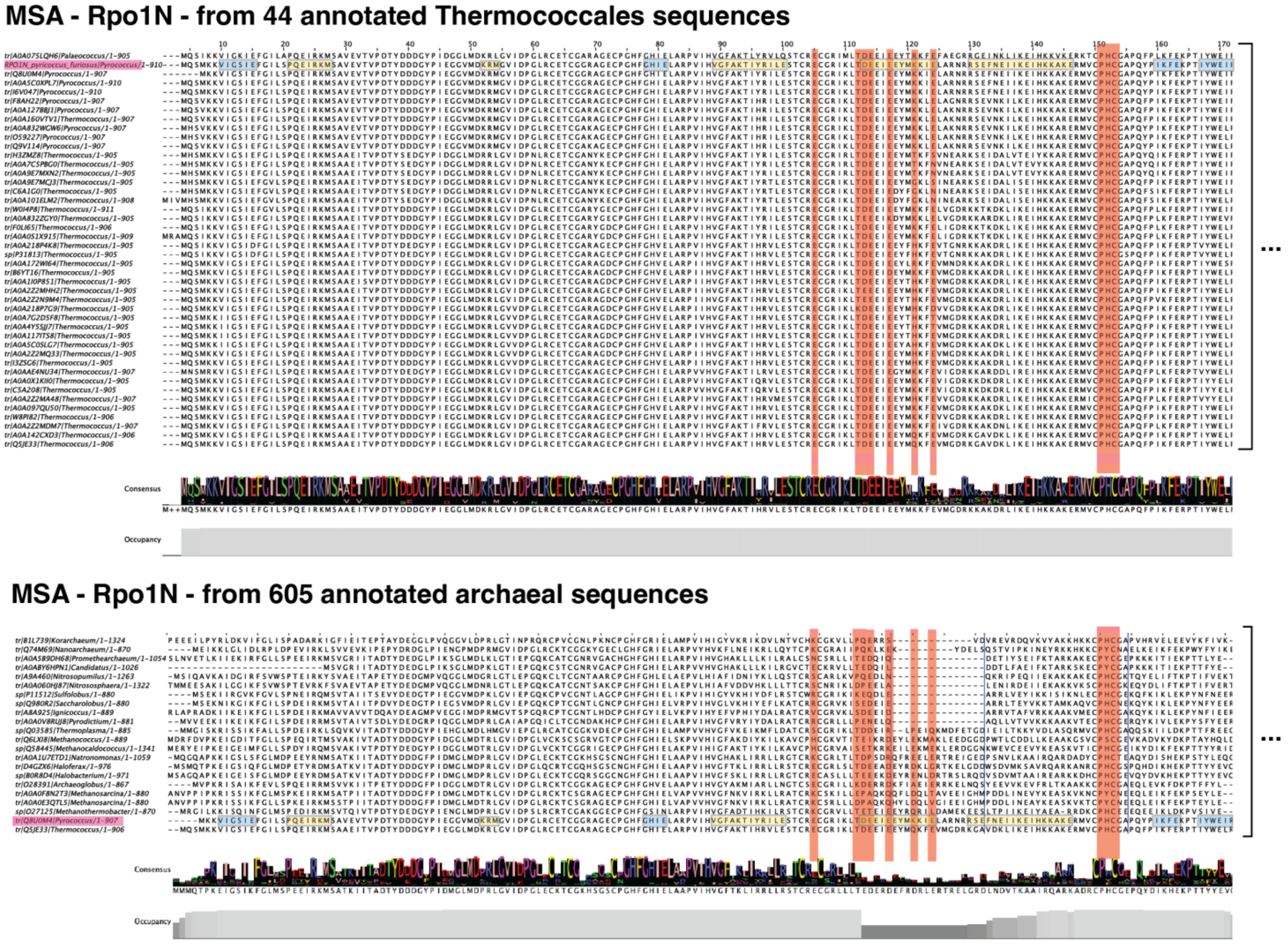


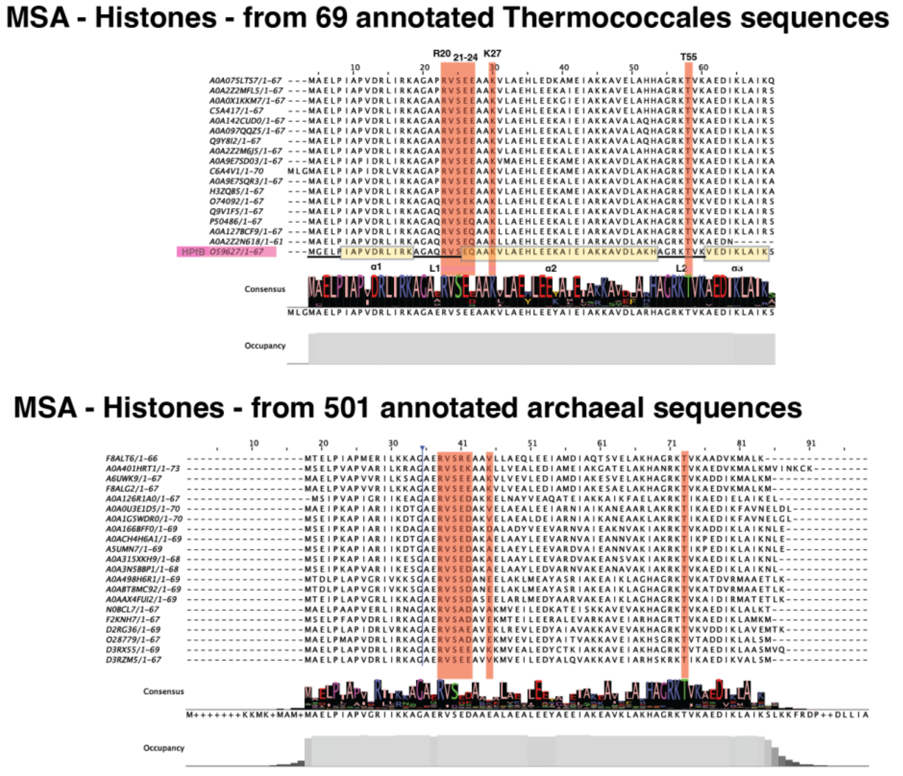


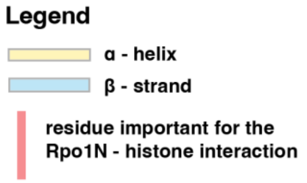


**Supplementary Figure 5. Conservation of Rpo1N and HPfB residues involved in the interaction interface.** Multiple sequence alignments of Rpo1N and HPfB sequences from *Thermococcales* and broader archaeal datasets. Alignments comprise Rpo1N sequences from 44 annotated *Thermococcales* and 605 annotated archaeal sequences, and histone sequences from 69 annotated *Thermococcales* and 501 annotated archaeal sequences. Residues identified as important for the Rpo1N-HPfB interaction are highlighted in red. Sequence conservation is represented by the consensus sequence and residue occupancy below each alignment. Figure generated using Jalview.

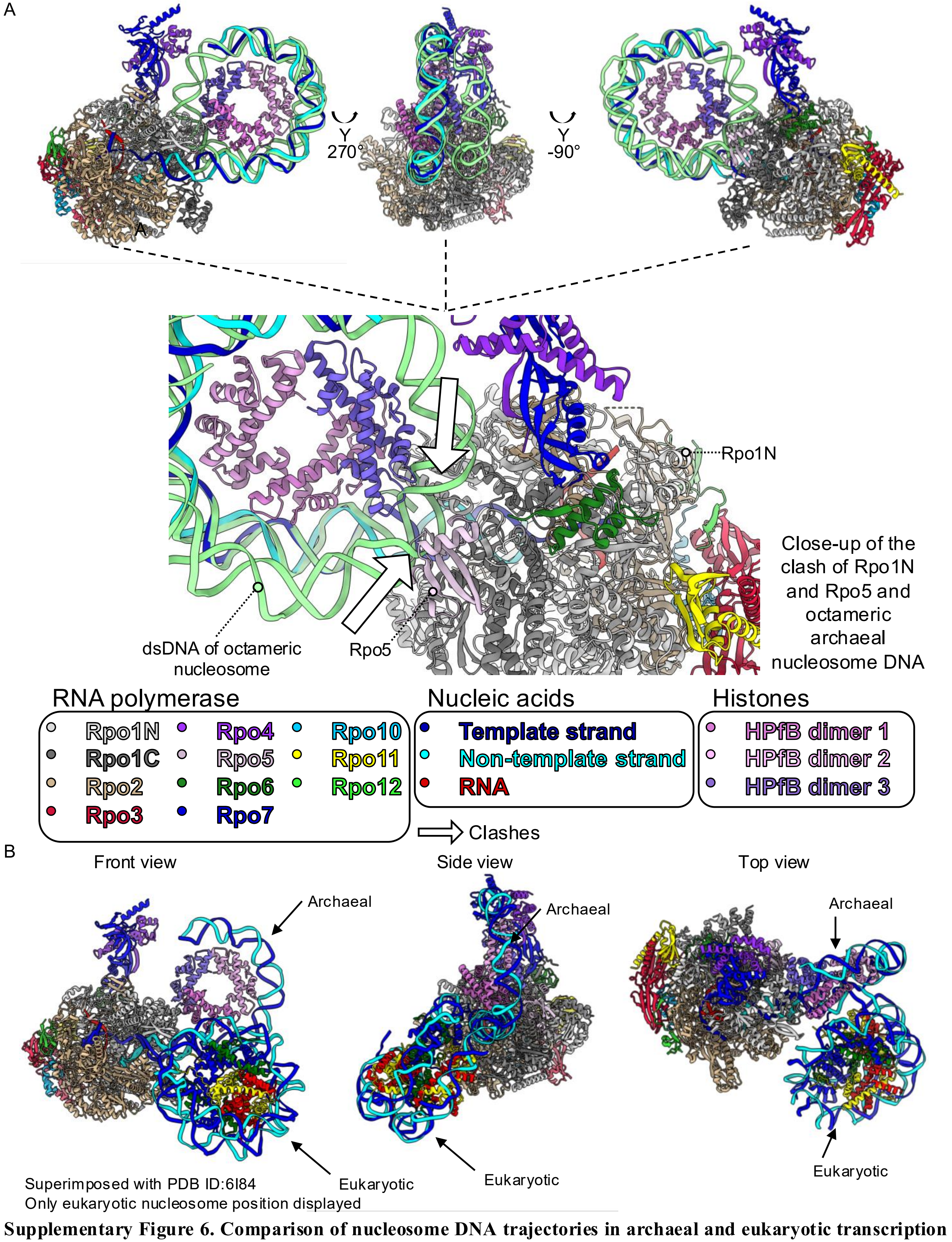


**Supplementary Figure 6. Comparison of nucleosome DNA trajectories in archaeal and eukaryotic transcription complexes.** A - Superimposition of the archaeal octasome structure (PDB ID: myoctasome) with the archaeal TEC-nucleosome complex. The DNA of the archaeal octasome is shown in green. Arrows indicate positions where the DNA trajectory of the archaeal octasome clashes with the Rpo1N and Rpo5 subunits of RNA polymerase. B - Superimposition of the archaeal TEC-nucleosome complex with a eukaryotic RNA polymerase II-nucleosome complex (PDB ID: 6I84). The structures are superimposed to compare the relative positioning of the nucleosome and the resulting DNA trajectories. The eukaryotic nucleosome is coloured according to the conventional colouring scheme.

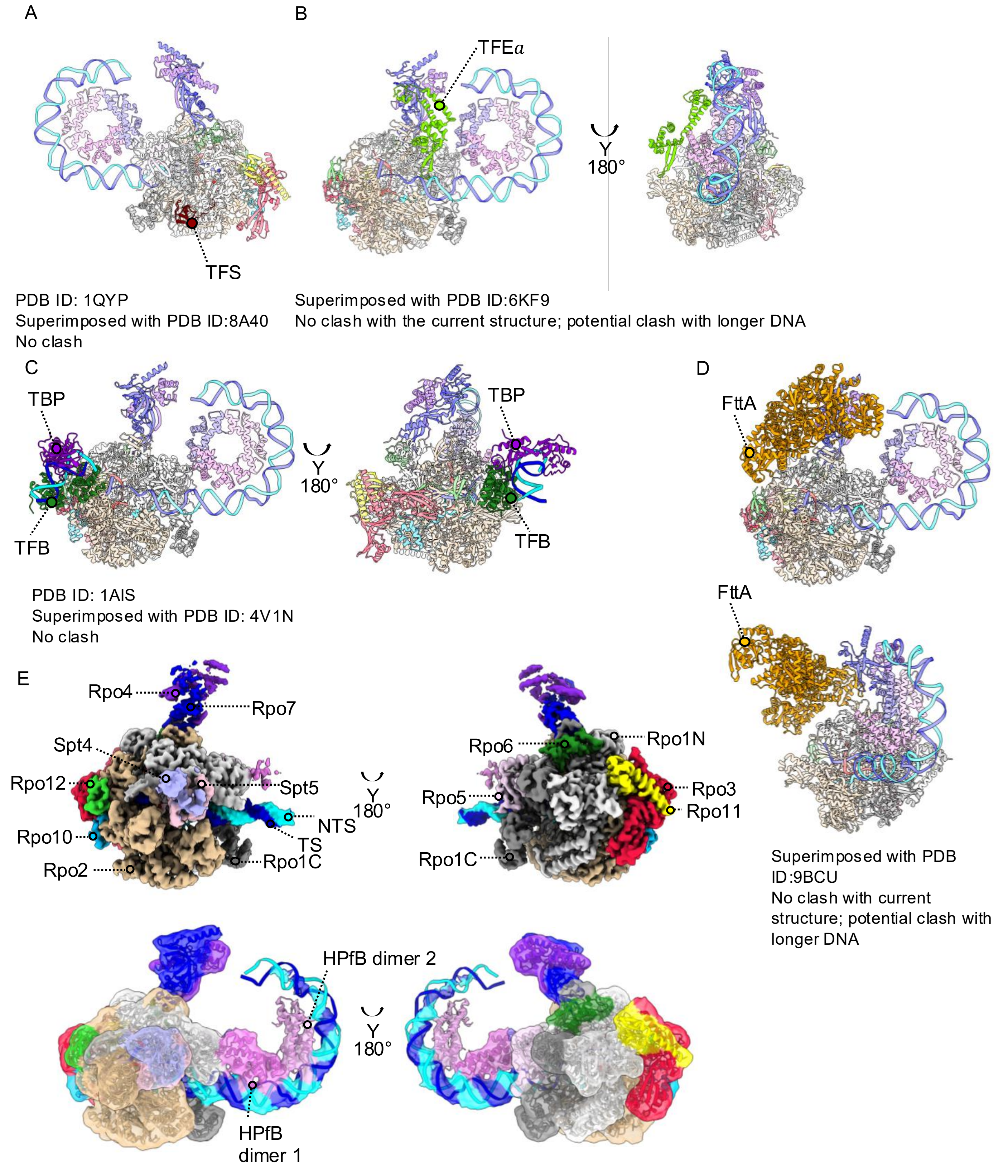


**Supplementary Figure 7. Structural compatibility of the archaeal TEC–nucleosome complex with transcription initiation, elongation and termination factors.** A - Superimposition of the archaeal TEC-nucleosome complex with the TFS (PDB 1QYP; superposed with PDB 8A40). No steric clash with the nucleosome is observed. B - Superimposition with the TFEα (PDB 6KF9). No clash is observed with the current DNA trajectory, although a potential steric clash may occur upon extension of the nucleosomal DNA. C - Superimposition with the TBP-TFB-DNA (PDB 1AIS; superimposed with PDB 4V1N). No steric clash with the nucleosome is observed. D - Superimposition with the FttA-containing complex (PDB 9BCU). No clash is observed with the current DNA trajectory, although a potential steric clash may occur upon extension of the nucleosomal DNA. E - Cryo-EM density map of the archaeal TEC-HPfB-Spt4/5 complex showing the positions of HPfB dimers relative to RNA polymerase. HPfB dimer 1 is clearly resolved, whereas density corresponding to HPfB dimer 2 is weaker and becomes apparent only at a higher contour level. RNA polymerase subunits, nucleic acids and HPfB dimers are coloured according to the scheme indicated previously.

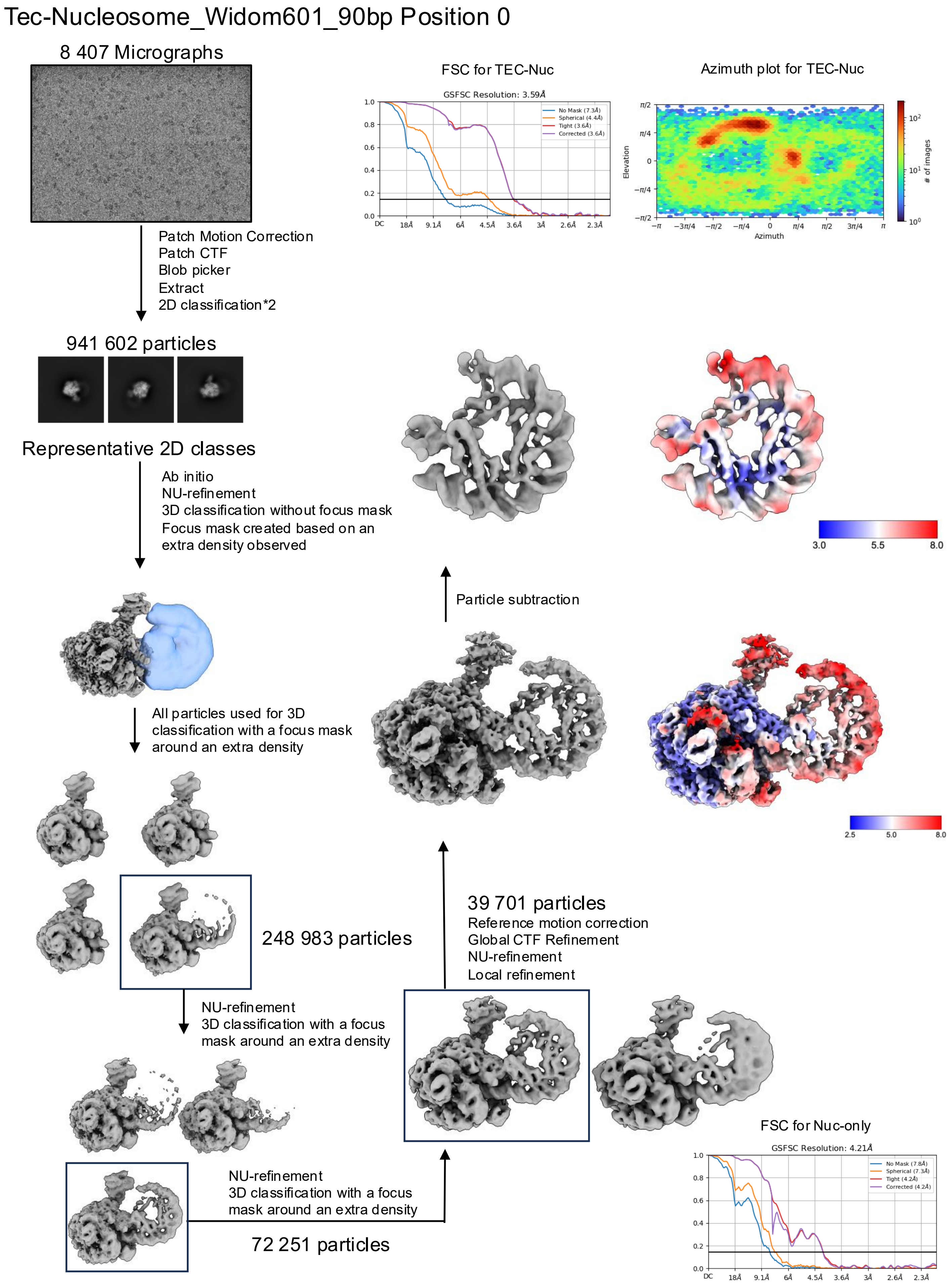


**Supplementary Figure 8. Cryo-EM data-processing workflow for the TEC–nucleosome complex at position 0.** The main data-processing steps are indicated. FSC curves, local resolutions, and an azimuth distribution plot for the final reconstructions are shown.

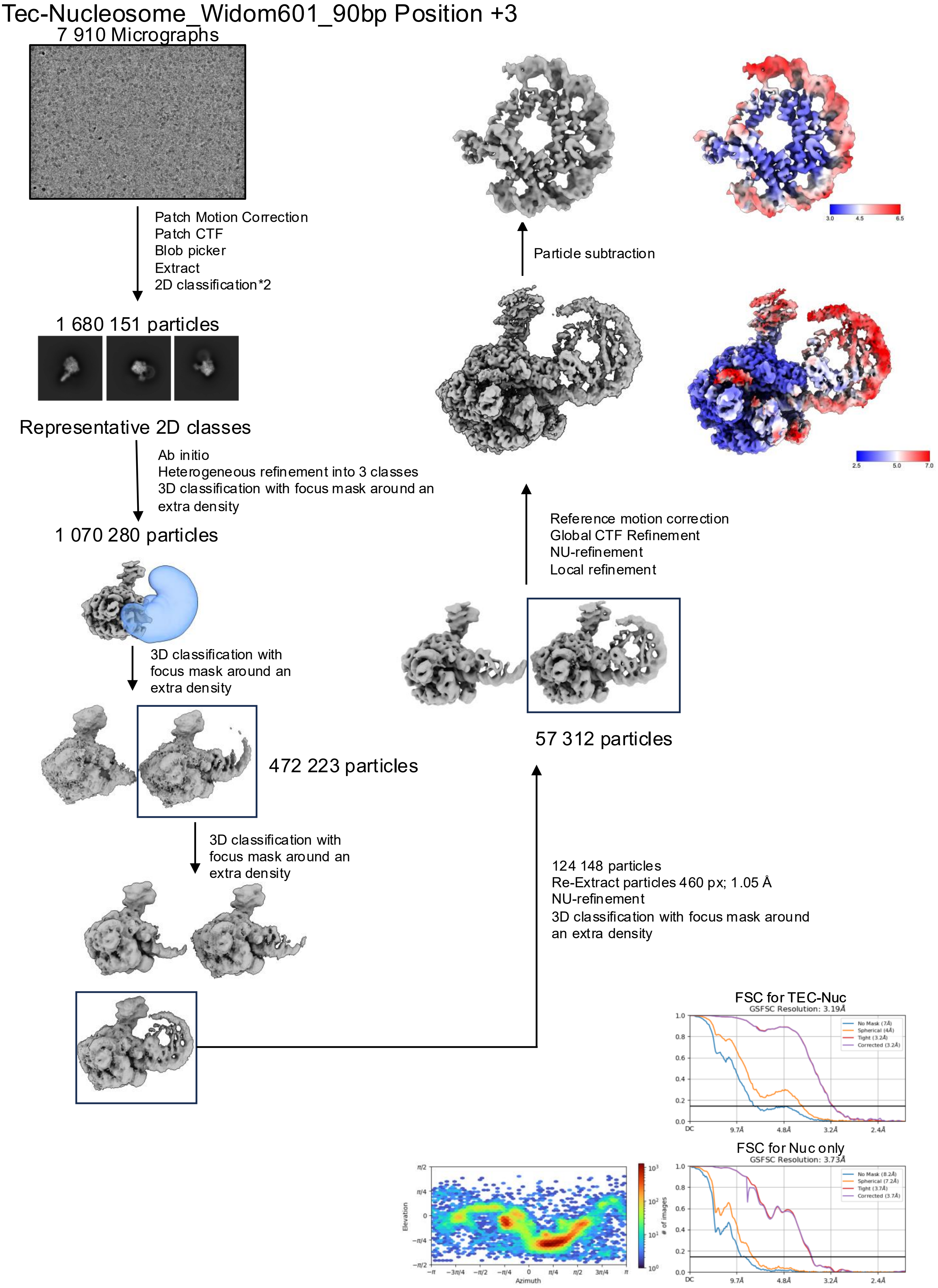


**Supplementary Figure 9. Cryo-EM data-processing workflow for the TEC-nucleosome complex at position +3.** The main data-processing steps are indicated. FSC curves, an azimuthal distribution plot and local and global resolution estimates for the final reconstructions are shown.

### Tec-Nucleosome_Widom601_90bp Position +6

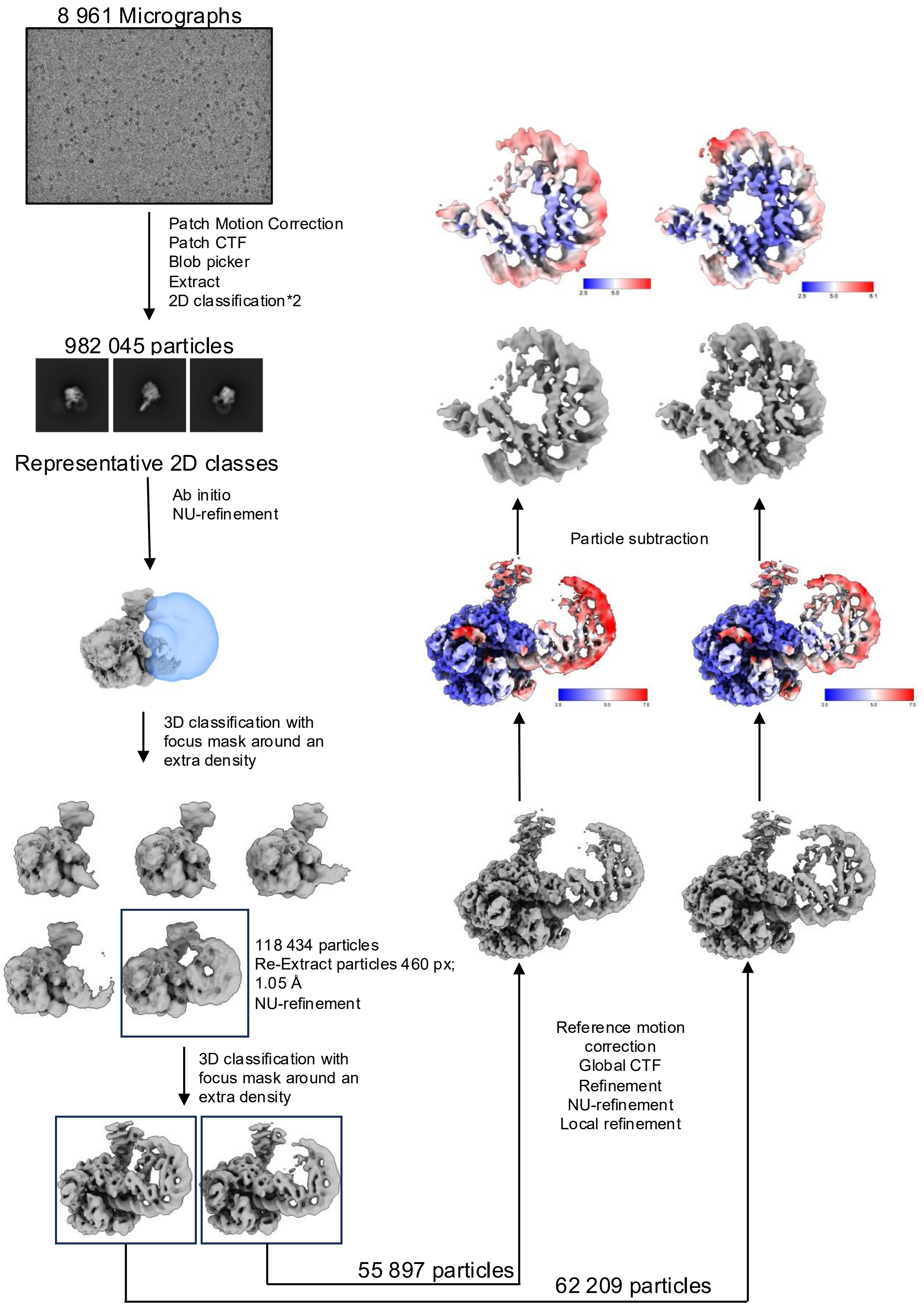


**Supplementary Figure 10. Cryo-EM data-processing workflow for the TEC-nucleosome complex at position +6.** The main data-processing steps are indicated. Local resolution maps for the final reconstructions are shown.

### Tec-Nucleosome_Widom601_90bp Position +6

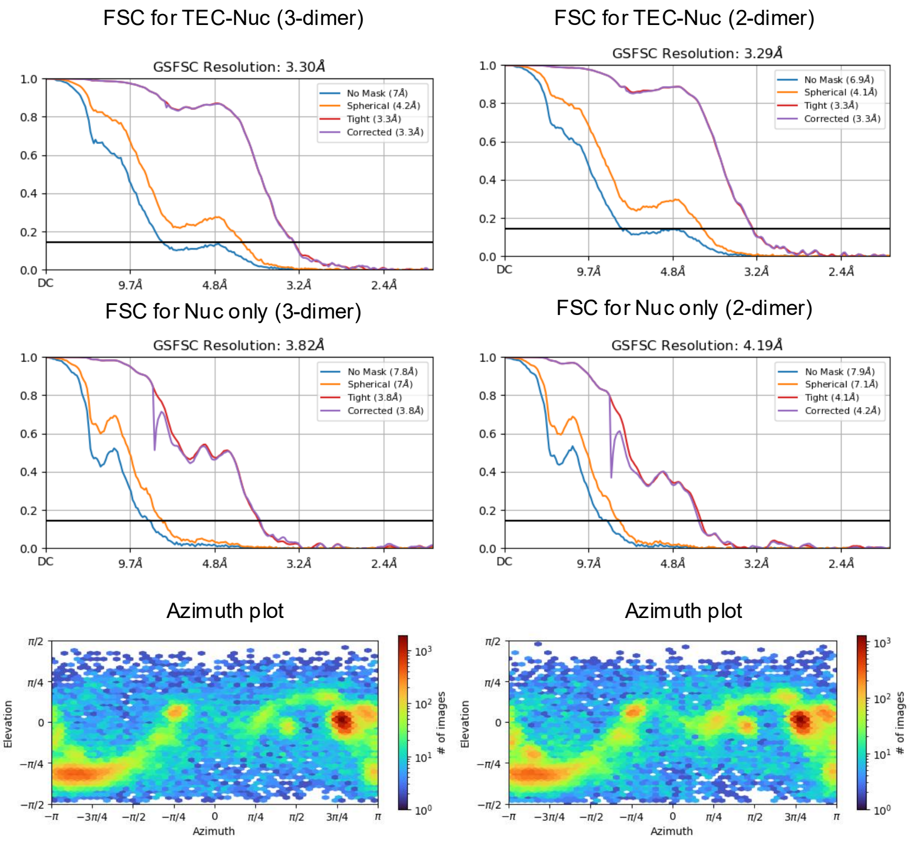


**Supplementary Figure 11. FSC and azimuth distribution analyses of the TEC-nucleosome complex at position +6.** FSC curves and azimuth distribution plots for the complete TEC-nucleosome complex and corresponding FSC nucleosome-only reconstructions are shown for the 3-dimer and 2-dimer HPfB states.

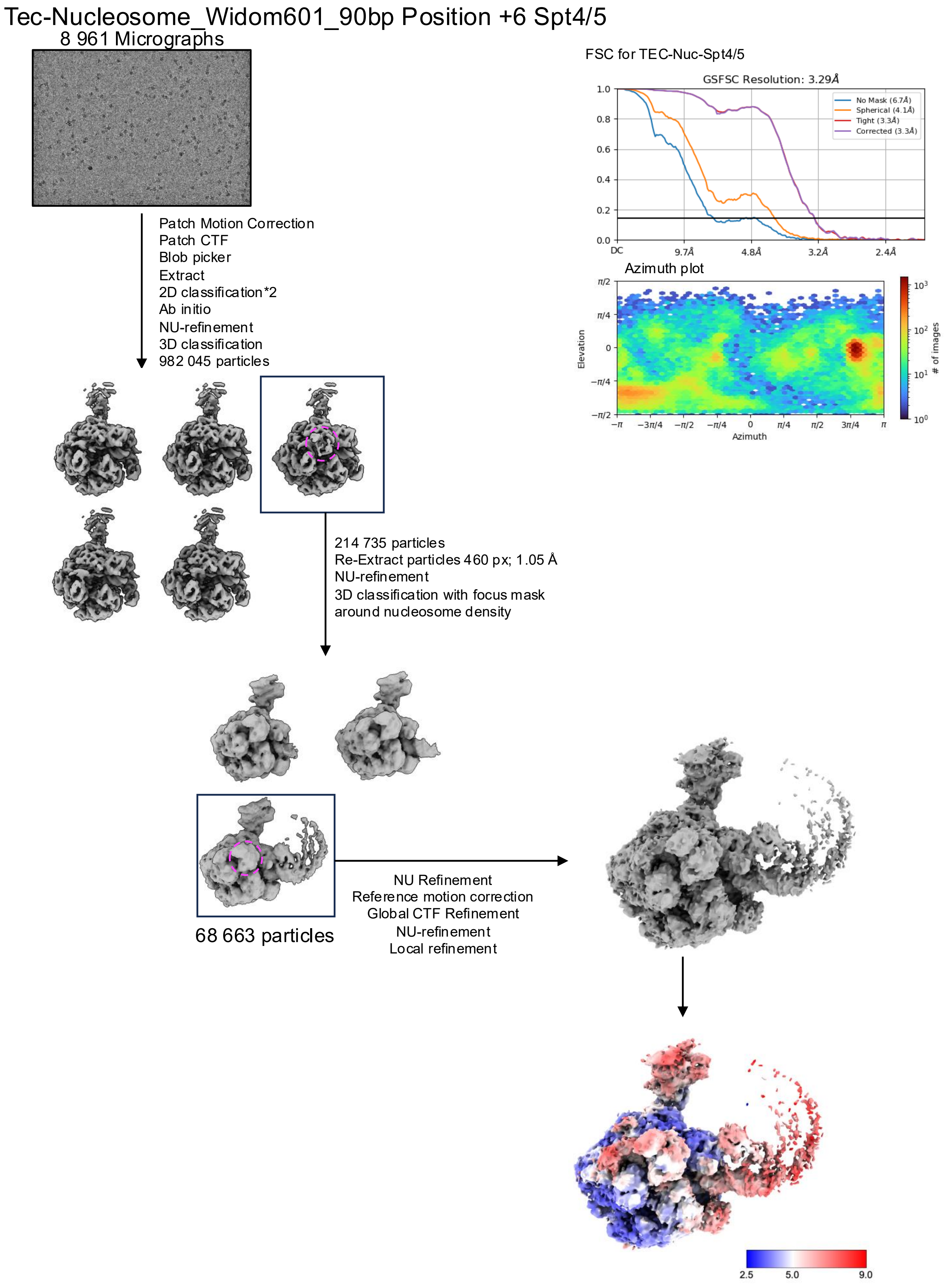


**Supplementary Figure 12. Cryo-EM data-processing workflow for the TEC-nucleosome–Spt4/5 complex at position +6.** The main data-processing steps are indicated. The FSC curve, azimuth distribution plot and local resolution map for the final reconstruction are shown.

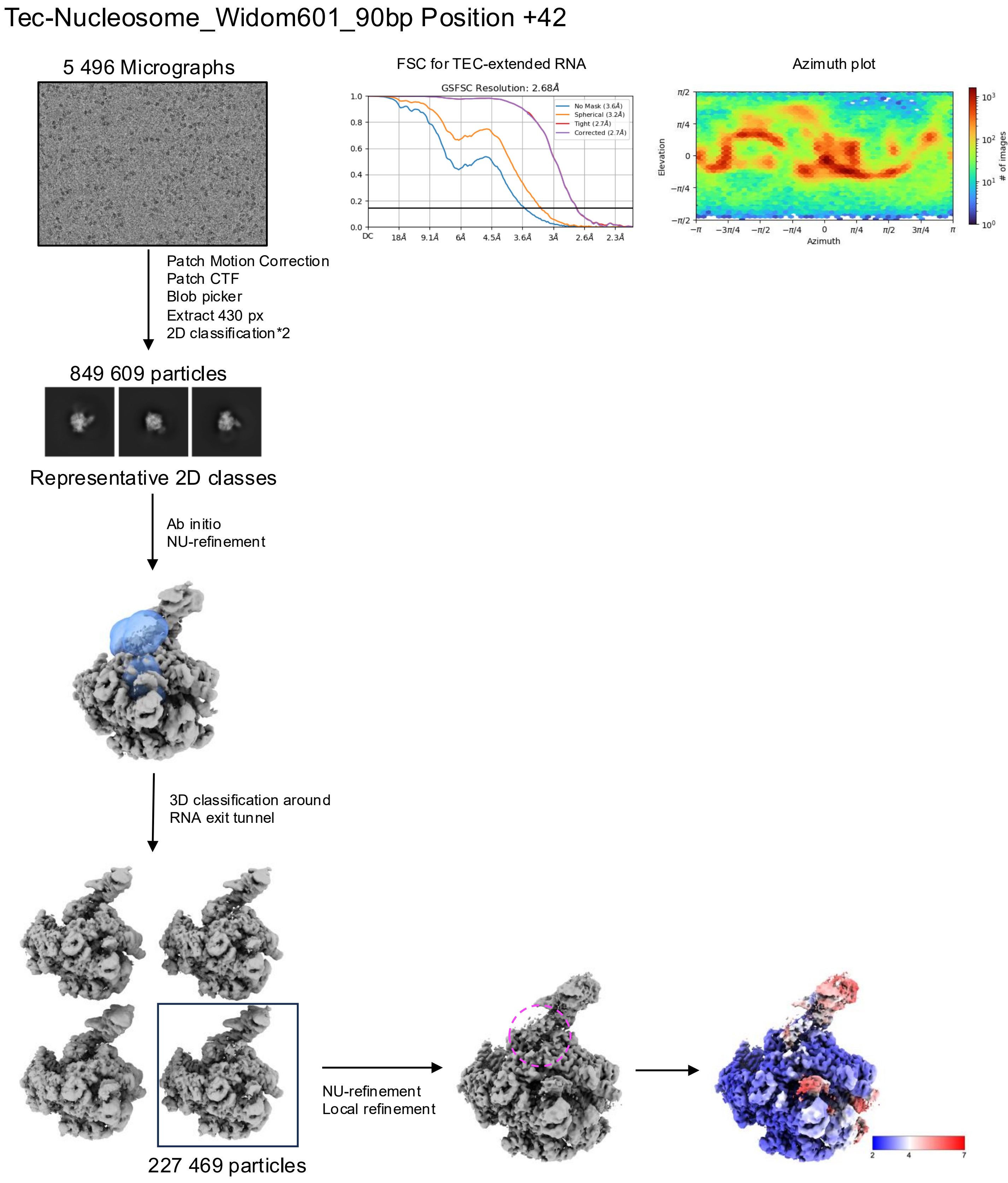


**Supplementary Figure 13. Cryo-EM data-processing workflow for the TEC complex at position +42.** The main data-processing steps are indicated. FSC and azimuth distribution plots, and a local resolution map for the final reconstruction are shown. HPfB based nucleosomewas not observed for this state.

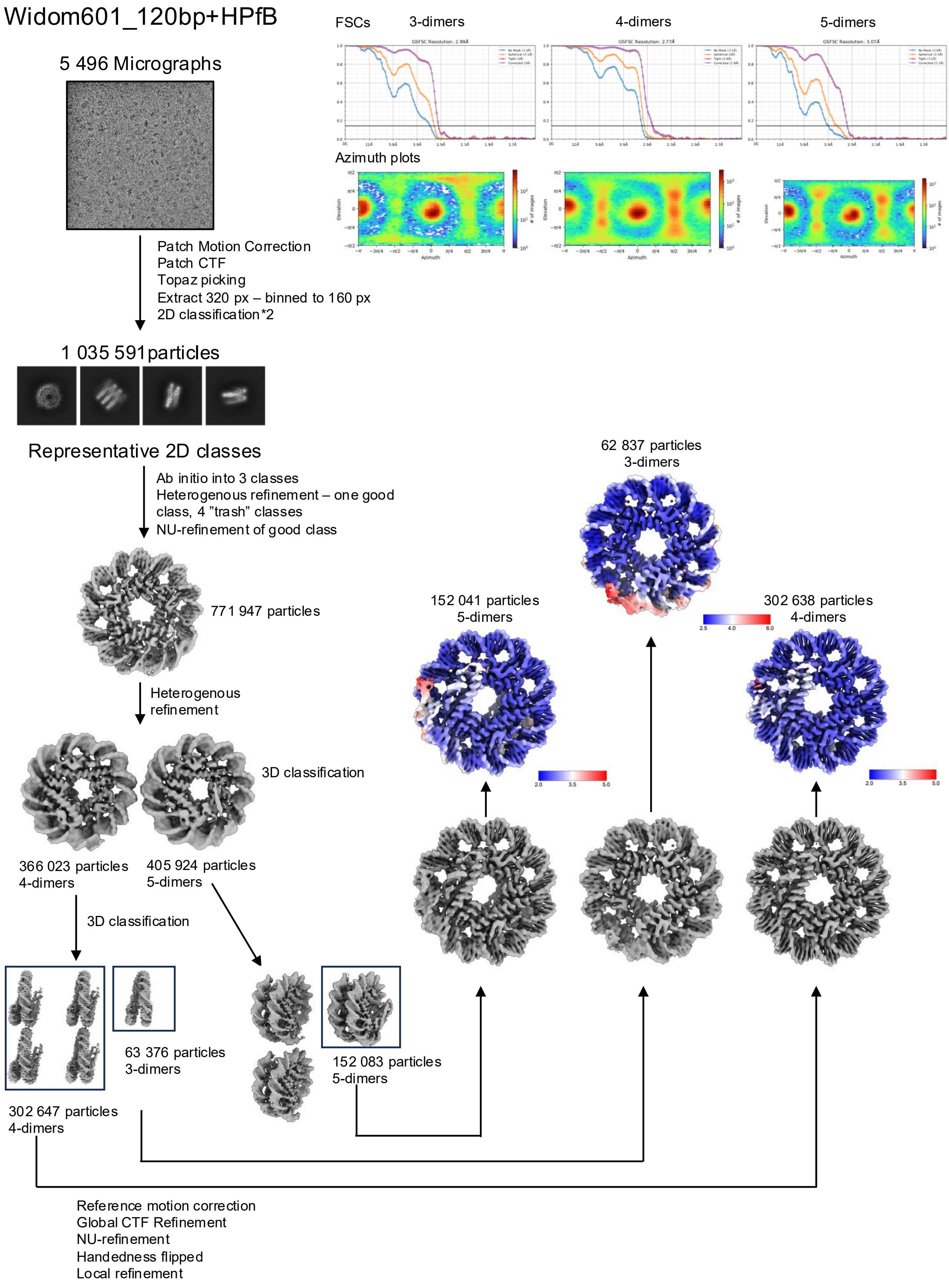


**Supplementary Figure 14. Cryo-EM data-processing workflow for the Widom601_120bp-HPfB nucleosomes.**
